# T-Cell Synaptosomes Orchestrate Long-Term Anti-Tumor Immunity via Proliferative and Metabolic Reprogramming

**DOI:** 10.1101/2025.11.06.687075

**Authors:** Sun-Kyoung Kang, Na-Young Kim, Sunghee Lee, Hyeonhee Lee, Won-Chang Soh, Jeong-Su Park, Hee-Tae Kang, Jihwan Park, Sunjae Lee, Yujeong Shim, Joonha Kwon, Hye-Ran Kim, Chang-Duk Jun

## Abstract

A defining feature of T-cell activation is its dependence on physical contact with antigen-presenting cells (APCs). During this interaction, activated T cells release microvesicles, known as T-cell immunological synaptosomes (TIS), onto the surface of APCs. While TIS are known to activate APCs, their broader *in vivo* functions remain largely unexplored. Here, we show that TIS are primarily phagocytosed by macrophages and dendritic cells, persist intracellularly for several days, and reprogram these cells in a manner distinct from LPS. TIS upregulate genes involved in metabolism, proliferation, and anti-inflammatory responses, and promote immune cell recruitment, including eosinophils. Notably, TIS induce minimal immune activation upon initial exposure but trigger a rapid and robust response upon secondary administration, indicating a priming effect reminiscent of adaptive immunity. In an MC38 colon cancer model, TIS treatment resulted in near-complete tumor suppression and durable protection upon rechallenge, highlighting their potential as a potent and long-lasting immunotherapeutic platform.

## INTRODUCTION

T-cell activation fundamentally depends on physical contact with antigen-presenting cells (APCs), where T-cell receptors (TCRs) engage peptide–MHC complexes at the immunological synapse (IS). Although this contact is primarily essential for antigen recognition, it is also thought to serve additional immunological functions, contributing to more complex layers of immune regulation. One such function is its involvement in promoting the release of extracellular vesicles (EVs). Notably, while soluble stimuli such as anti-CD3/CD28 antibodies or PMA/ionomycin efficiently trigger T-cell activation and TCR internalization ^1–3^, they fail to elicit detectable EV release ^4^. In contrast, T cells stimulated by immobilized ligands or peptide-presenting APCs produce substantial amounts of EVs, highlighting the strict dependence of vesicle biogenesis on adhesion-mediated signaling, particularly via the ICAM-1/LFA-1 axis ^4,5^.

Given that these EVs are released in small quantities from T cells and selectively adhere to the surface of APCs, their precise function remains elusive. One might intuitively assume that they mirror T-cell activity, yet their limited, antigen-specific release and restricted delivery to APCs *in vivo* suggest that their physiological effects may be confined to adaptive immune responses. This raises a critical question: what would happen if these T cell-derived vesicles were harvested in bulk *ex vivo* and administered systemically? Would they simply replicate a portion of adaptive T-cell function, or could they exert broader influence across the immune system? At present, little is known about their capacity to modulate immune responses beyond the narrow context of physiological antigen recognition. In contrast to exosomes and other EVs, which are generally regarded as mediators of long-range intercellular communication ^6,7^, TIS may not only echo T-cell activity but also reveal antigen-independent functions that could redefine the boundaries of adaptive immunity.

Despite increasing interest in the immunological potential of T cell-derived EVs, their precise origin and mode of biogenesis remain subjects of active investigation. Early studies often referred to these vesicles as exosomes, despite the lack of direct morphological evidence supporting endosomal origin ^7–10^. More recent reports instead describe their release as a form of ectocytosis occurring at the IS, wherein vesicles bud from the central supramolecular activation cluster (c-SMAC) via ESCRT-dependent membrane scission ^11–13^. However, given that ectocytosis has classically been defined as an adhesion-independent process ^14^, this mode of release appears mechanistically distinct from the adhesion-dependent vesicle production observed during physiological T-cell activation. Intriguingly, ESCRT components are also localized to the distal tips of T-cell microvilli, where they mediate membrane fragmentation during the contraction or kinapse phase that follows IS formation ^5^. This dual localization raises the possibility that synaptic ectosomes ^11–13^ and microvilli-derived vesicles—termed T-cell microvilli particles (TMPs) ^4,5^—may in fact represent a shared biological entity ^15–17^. Unlike synaptic ectosomes, which have been primarily described as c-SMAC–restricted structures, TMPs are generated more broadly through microvilli fragmentation ^5^. Nonetheless, considering that the trailing edge of migrating T cells recapitulates features of the c-SMAC, it is plausible that TMPs and ectosomes originate from a common biogenetic program. For this reason, we collectively refer to these vesicles as T-cell immunological synaptosomes (TIS), a term that emphasizes both their synaptic origin and their selective communication with APCs.

Intriguingly, systemic administration of TIS led to their rapid uptake and prolonged persistence in macrophages and dendritic cells (DCs). Unlike classical TLR ligands such as LPS, TIS did not provoke immediate inflammation but primed APCs for robust recall responses, resulting in potent tumor suppression and durable protection upon rechallenge. Mechanistically, TIS induced proliferating macrophage clusters and recruited eosinophils, both essential for their antitumor effects. These delayed yet amplified responses parallel the concept of trained innate immunity, suggesting that TIS are not merely mimics of T-cell activity but potent inducers of innate reprogramming that redefine the interface between adaptive and innate immunity—and establish a vesicle-based platform for durable cancer immunotherapy.

## RESULTS

### TIS are adhesion-dependent vesicles that remain functionally stable and activate professional APCs

As shown in Figure 1A, TIS arise from finger-like microvillar projections of activated T cells, appearing as rod-shaped extensions that become spherical vesicles upon adhesion. For *in vivo* studies, we generated TIS with anti-CD3/CD28–coated beads ^18^, producing mainly round vesicles enriched in TCRζ and ARRDC1 (Figure S1A). Their release was adhesion-dependent, as soluble antibodies induced minimal production (Figure S1B). Functionally, TIS from CD3⁺, CD4⁺, or CD8 ⁺ T cells activated DCs and macrophages, upregulating MHC II, nitric oxide, and TNF-α (Figures 1B and S1C). Activity was maintained after freeze–thaw or lyophilization, and TIS potentiated LPS-induced nitric oxide and TNF-α, indicating that they amplify classical innate pathways. Because CD4⁺ T cells form stable synapses with DCs, we focused on CD4⁺ T cell–derived TIS (4-TIS) for subsequent experiments.

**Figure 1.**
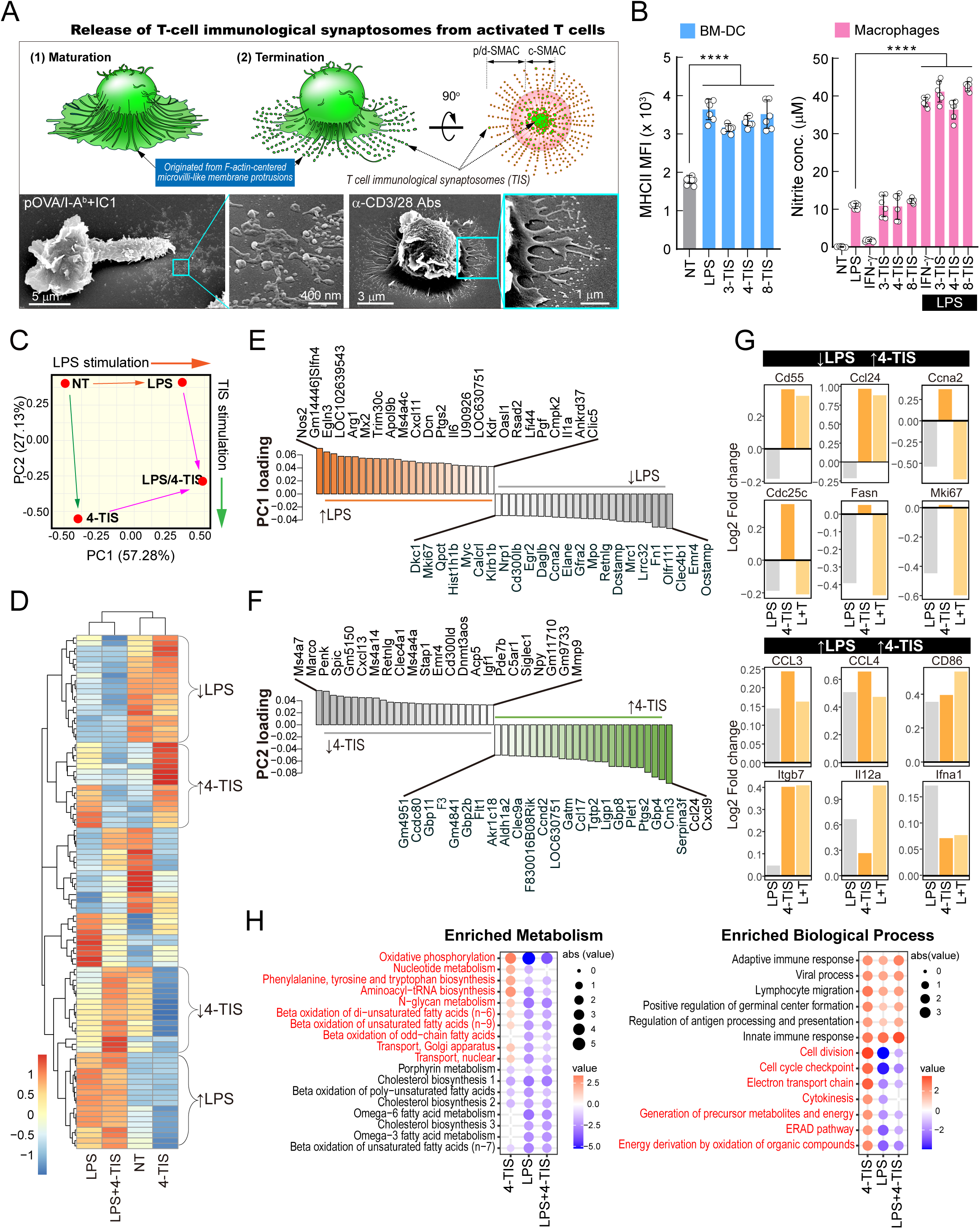
4-TIS elicit distinct gene signatures and adaptive-like immune modulation compared to LPS-driven inflammatory programs. (A) Schematic of TIS release during IS maturation and termination. Representative SEM images of naïve OT-II CD4⁺ T cells stimulated with immobilized OVA/I-Aᵇ + ICAM-1 or anti-CD3/CD28, highlighting microvilli-like protrusions and TIS release sites (Cyan boxes). (B) Surface MHC II expression on BMDCs (left) and nitric oxide production by macrophages (right) after 24 h stimulation with NT, LPS, or TIS derived from CD3⁺, CD4⁺, or CD8⁺ T cells. Data represent mean ± SD of three independent experiments; ****p < 0.0001 versus NT (left) or LPS (right), one-way ANOVA with Tukey’s correction. (C) PCA of transcriptomes from NT-, LPS-, 4-TIS–, and LPS+4-TIS–treated BMCs, with arrows indicating transcriptional trajectories. (D) Heatmap of differentially expressed genes (rows) across treatments (columns). (E, F) PCA gene loadings highlighting LPS-driven (E) and TIS-driven (F) programs. (G) Log2 fold changes of representative genes suppressed by LPS but induced by TIS (top), or induced by both (bottom). (H) Functional enrichment analysis of DEGs. Left, KEGG metabolic pathways; right, GO biological processes. Dot color represents normalized enrichment score (NES; red, positive; blue, negative), dot size reflects absolute NES.

### Transcriptomic profiling reveals that 4-TIS reprogram myeloid cells toward adaptive-like, non-inflammatory gene programs

To assess their impact, GM-CSF–differentiated bone marrow cells (BMCs) were treated with TIS, LPS, or both and analyzed using Affymetrix Mouse Gene ST 2.0 arrays. PCA and heatmap clustering revealed that TIS induced a transcriptional profile distinct from the classical inflammatory program triggered by LPS (Figures 1C and 1D). Differential expression showed that TIS upregulated genes involved in immune regulation and tissue remodeling (*Ptgs2, Clec9a, Aldh1a2*) and interferon-associated “trained immunity” signatures (*Gbp2b, Gbp8, Gbp11*), while suppressing pro-inflammatory and tissue-destructive genes (*Mmp9, Marco, Siglec1, C5ar1*) (Figures 1E and 1F). By contrast, LPS strongly induced canonical TLR4-driven inflammatory responses, including interferon-stimulated genes (*Ifit, Rsad2, Oasl1*) and cytokines (*Il6, Il1b*), while repressing reparative programs (*Mrc1, Egr2, Fn1*) (Figure 1G). Pathway analysis indicated that TIS enhanced oxidative phosphorylation, amino acid metabolism, and energy transport, while reducing cholesterol synthesis and fatty acid oxidation (Figure 1H). Consistently, GO enrichment highlighted cell cycle, metabolic, and adaptive immune–associated pathways in TIS-treated cells, in contrast to the acute inflammatory response dominated by LPS (Figure 1H).

### 4-TIS orchestrate adaptive-like cytokine programs through T cell–derived factors

To assess the contribution of T cell–derived factors to TIS-mediated immune modulation, we examined their cytokine composition and effects on APCs. Cytometric bead array revealed that TIS, particularly from activated CD4⁺ and CD8⁺ T cells, contained multiple cytokines, including IFN-γ, MIP-2 (CXCL2), MIP-1α (CCL3), TNF-α, GM-CSF, IL-17A, and XCL1, reflecting selective acquisition of T cell effector molecules (Figure S2A). Functionally, TIS-stimulated BMDCs and macrophages preferentially induced cytokines associated with adaptive immunity (CCL24, CXCL9, IFN-γ, CCL22, CCL7), whereas LPS predominantly triggered innate inflammatory cytokines (CXCL1, CXCL10, TNF-α, IL-6) (Figures S2B and S3). Shared cytokines such as CCL2 and CCL3 were induced by both stimuli. Given the abundance of IFN-γ in TIS, we tested its role using TIS from IFN-γ– deficient T cells. These vesicles induced markedly lower levels of IFN-γ, CXCL9, CXCL10, and TNF-α in DCs (Figure S2C), establishing IFN-γ as central to TIS-triggered immune modulation. Blockade of TNF-α and GM-CSF, alone or in combination, further reduced DC cytokine output, supporting a cooperative network of T cell–derived cytokines in orchestrating the adaptive-like immune responses induced by TIS (Figure S2D).

### 4-TIS promote delayed but broad recruitment of immune cells in vivo

Given the unique gene and cytokine programs induced by TIS *ex vivo*, we next asked whether these features translate into a coordinated immune landscape *in vivo*. As shown in the experimental timeline (Figure 2A), peritoneal cells were collected at multiple time points after injection. Unlike LPS or poly(I:C), which induced a rapid but transient influx of leukocytes, TIS triggered a delayed yet robust accumulation of diverse immune populations (Figure 2B). Total peritoneal cell numbers increased by day 8 and peaked on day 11, with the CD45⁺ compartment enriched for CD4⁺ and CD8⁺ T cells, CD19⁺ B cells, MHCII⁺ DCs, and Siglec-F⁺ eosinophils (Figures 2C and S4). In contrast, LPS rapidly recruited CD11b⁺Ly6G⁺ neutrophils within 4 h, whereas TIS elicited only a mild and delayed neutrophil response (Figure 2C). By day 11, both CD4⁺ and CD8⁺ T cells expanded, alongside significant increases in DCs and macrophages (Figures 2C and S4). Macrophages expanded in both LPM and SPM subsets, in contrast to the LPS-driven reduction in LPMs (Figure 2C). A distinct SSC^high^ Siglec-F⁺ eosinophil population also emerged, likely supported by *Ccl24* induction in GM-CSF–differentiated BMCs (Figures 1F and 1G).

**Figure 2.**
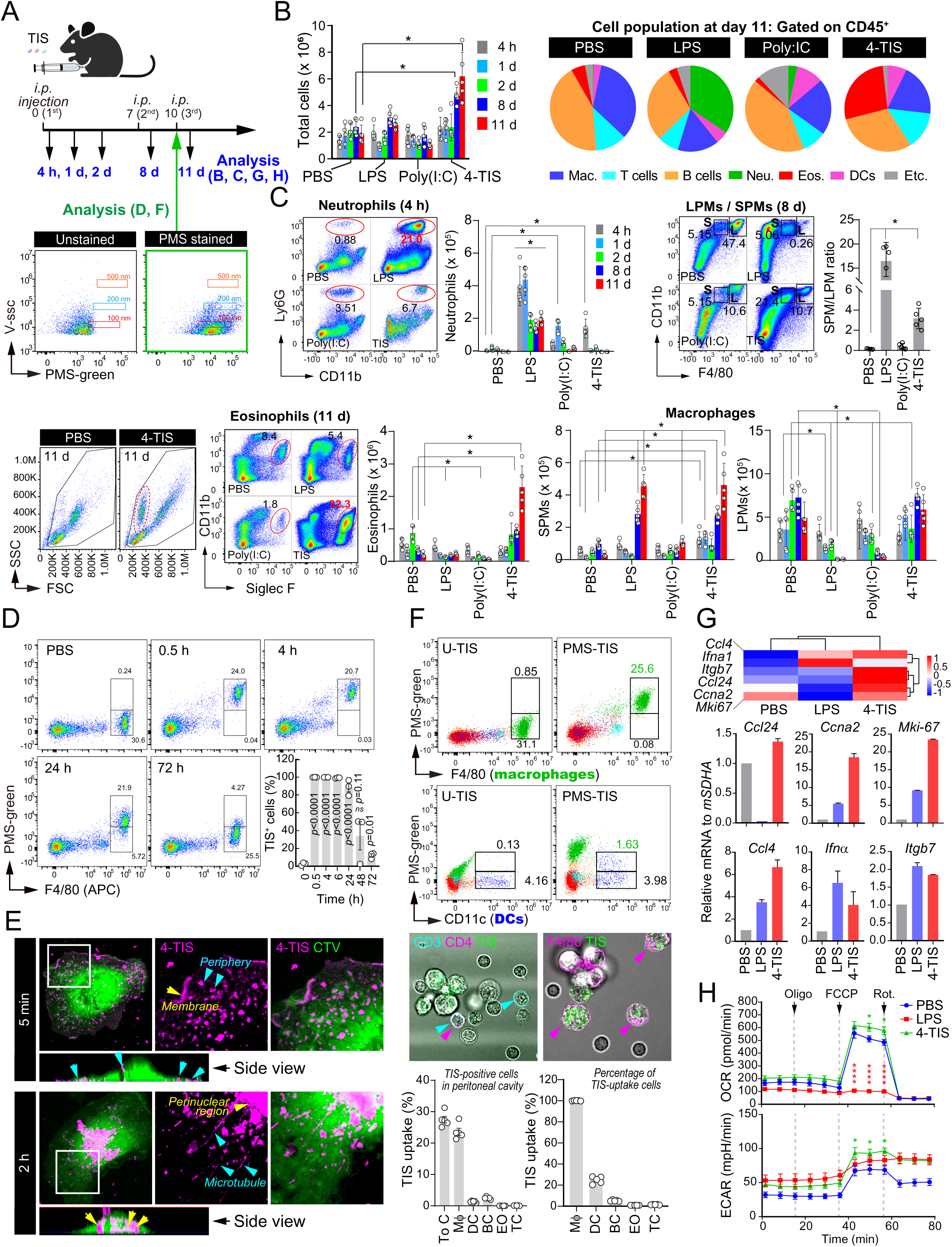
Unlike LPS, 4-TIS induce delayed but broad immune recruitment with adaptive-like features in the peritoneum. (A) Experimental design for i.p. injection of PBS, LPS (25 μg), Poly(I:C) (50 μg), or 4-TIS, and peritoneal cell collection at the indicated time points. Representative flow cytometry plots of PMS-green–labeled TIS (size calibrated with 100–500 nm beads). (B) Total peritoneal cell counts at 4 h, 1 d, 2 d, 8 d, and 11 d (left). Pie charts show immune composition of CD45⁺ cells at day 11 (right). Data represent mean ± SEM; n = 4–5 mice per group; *p* < 0.05, one-way ANOVA with Tukey’s test. (C) Quantification of neutrophils, eosinophils, and macrophages (further subdivided into LPMs and SPMs) at indicated time points. Data represent mean ± SEM; n = 4–5 mice per group; *p* < 0.05, two-way ANOVA. (D) Uptake kinetics of PMS-labeled TIS in F4/80⁺ macrophages by flow cytometry. Percentages of TIS⁺ macrophages are plotted over time. (E) Confocal imaging of macrophages exposed to fluorescently labeled TIS. TIS localized to the plasma membrane at 5 min (arrowheads) and to perinuclear microtubules at 2 h. (F) Uptake of PMS-labeled TIS 4 h after the third injection. Left, proportion of PMS⁺ cells among peritoneal leukocytes. Right, uptake within subsets (macrophages, DCs, B cells, eosinophils, T cells). Data represent mean ± SEM; n = 3 mice per group. (G) Validation of microarray findings. Heatmap of representative transcripts. RT-qPCR (See Fig. 1G) confirmed upregulation of *Ccl24, Ccl4, Ccna2, Mki67, Ifna1*, and *Itgb7* in macrophages after repeated TIS injections. Data represent mean ± SD from three independent experiments; *p* < 0.05, Student’s *t*-test. (H) Seahorse analysis of peritoneal macrophages stimulated with PBS, LPS, or TIS. Oxygen consumption rate (OCR, top) and extracellular acidification rate (ECAR, bottom) are shown. Data represent mean ± SEM of technical triplicates; representative of three independent experiments.

To test whether this recruitment pattern was confined to the peritoneum, we examined intradermal TIS administration. Unlike LPS or poly(I:C), which induced early neutrophil-dominant infiltration, TIS drove a delayed accumulation of macrophages, lymphocytes, and eosinophils, accompanied by increased skin thickness and leukocyte counts on days 8–11 (Figures S6A and S6B). TIS also upregulated chemokines such as CCL24, CCL9, and CCL7 in skin and serum (Figure S6C), indicating that TIS promote coordinated recruitment of innate and adaptive immune cells across peripheral tissues.

### Early in vivo tracking revealed rapid uptake of TIS by professional APCs

To examine TIS uptake kinetics, fluorescently labeled TIS (PMS Green) were injected intraperitoneally (Figure 2A). Flow cytometry showed that peritoneal macrophages (F4/80⁺ CD11b⁺) rapidly internalized TIS within 0.5 h, peaking at 4 h and persisting up to 72 h (Figure 2D). DCs also displayed appreciable uptake, B cells exhibited transient uptake that declined within 24 h, eosinophils internalized only small amounts, and T cells remained largely negative (Figure S5). After repeated dosing, macrophages and DCs remained the dominant TIS-positive populations, whereas B- and T-cell uptake was minimal (Figure 2F). Confocal imaging confirmed that within 5 min TIS localized along the plasma membrane or appeared as puncta at the periphery, and by 2 h aligned with microtubules and accumulated near the perinuclear microtubule-organizing center, consistent with trafficking to phagolysosomes (Figure 2E).

Transcriptomic profiling of peritoneal macrophages isolated after repeated *in vivo* TIS administration revealed gene expression patterns closely resembling those induced by TIS-treated BMCs (*Ccna2*, *Mki67*, *Ccl24*, *Ccl4*, *Itgb7*, *Ifna1*) (Figure 1G and Figure 2G, top). These changes were validated by RT-qPCR (Figure 2G, bottom), confirming that macrophages adopt a proliferative and immune-stimulatory phenotype in response to TIS *in vivo*. Functionally, TIS-treated macrophages exhibited increased oxygen consumption rate (OCR) compared with PBS and LPS, indicating enhanced oxidative phosphorylation (Figure 2H). Extracellular acidification rate (ECAR) was elevated relative to PBS and comparable to that induced by LPS, suggesting that TIS also engages glycolysis (Figure 2H). Thus, TIS reprogram macrophages into a metabolically active state characterized by combined engagement of oxidative phosphorylation and glycolysis, distinct from the glycolysis-dominant program induced by LPS. Notably, TIS also increased OCR and ECAR in B cells (Figure S7), demonstrating that their metabolic effects extend beyond macrophages. Together, these findings identify peritoneal macrophages and DCs as primary responders to TIS, mediating uptake, intracellular trafficking, and downstream immune activation.

### 4-TIS preferentially reprograms peritoneal macrophages and drives a proliferative eosinophil response

To define the cellular programs induced by TIS, we performed single-cell RNA sequencing of peritoneal cells from PBS-, LPS-, or TIS-treated mice. UMAP clustering revealed that TIS markedly remodeled the peritoneal immune landscape, most prominently within the monocyte–macrophage compartment (Figures 3A–3C) ^20^. A proliferating macrophage (PM) cluster emerged exclusively in TIS-treated mice, enriched for cell-cycle genes such as *Mki67*, *Top2a*, and *Birc5* (Figures 3C, 3G, and S9B). Canonical markers confirmed cluster identities (Figure S8), and pathway analyses indicated that TIS-associated macrophages adopted a metabolically active, regulatory phenotype distinct from the acute inflammatory profile induced by LPS (Figures 3D–3F). In addition, TIS-stimulated macrophages selectively upregulated genes including *Slc7a2*, *Rnase2a*, *Chil3*, *Ucp1*, *Kcnn3*, *Cldn10*, *Clec4d*, and *Pilra*, highlighting their acquisition of a unique transcriptional program (Figure S9A).

**Figure 3.**
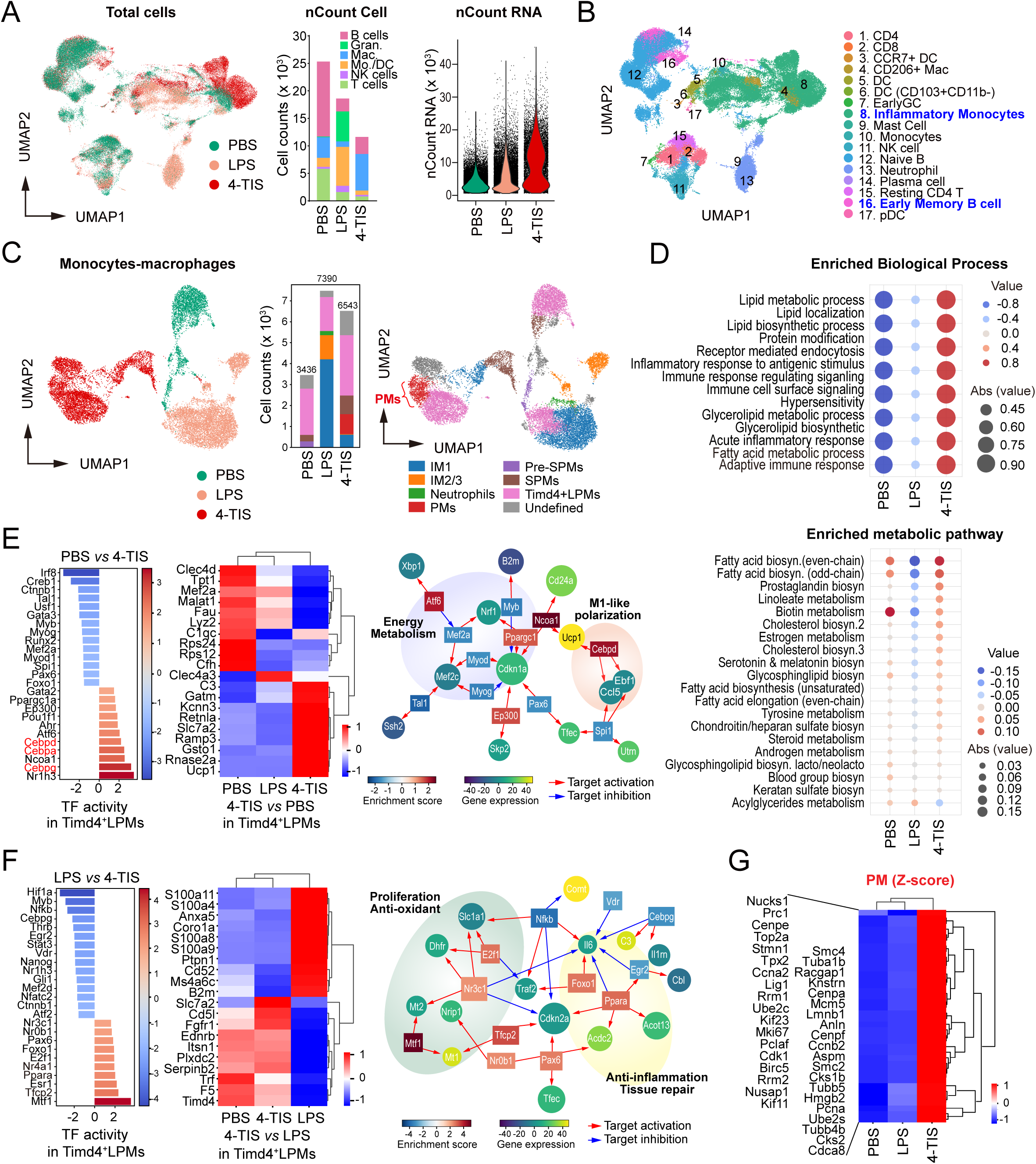
Single-cell analysis revealed that 4-TIS primarily reprogram peritoneal macrophages into proliferative and metabolically active states. (A) UMAP of all peritoneal cells from PBS-, LPS-, and 4-TIS–treated mice, colored by treatment. Bar graphs show cell numbers (nCount) per subset; violin plots display total RNA counts per cell. (B) UMAP colored by immune cell annotations. Numbers denote cluster IDs; clusters 8 (inflammatory monocytes) and 16 (early memory B cells) were selectively altered by 4-TIS. (C) Macrophage-focused analysis. Left: UMAP colored by treatment. Middle: stacked bar plots showing relative proportions of macrophage subtypes across treatments. Right: pooled UMAP annotated as pre-SPMs, SPMs, LPMs, IM1–3, neutrophils, and proliferating macrophages (PMs). (D) GO biological processes and KEGG metabolic pathways enriched in 4-TIS versus PBS/LPS. Dot color represents enrichment score (red, up; blue, down); dot size indicates absolute value. (E, F) Transcriptional reprogramming of Timd4⁺ LPMs. (E) Differential activity in 4-TIS versus PBS; (F) 4-TIS versus LPS. Left: transcription factor (TF) activity scores. Middle: representative gene heatmaps (row z-scores). Right: TF–target networks (red, activation; blue, inhibition). (G) Heatmap of proliferation-related genes (*Mki67, Top2a, Birc5, Ccnb2, Cenpf*) showing selective induction in 4-TIS–treated macrophages. Data represent mean ± SEM; n = 3 mice per group. Differential expression tested by Wilcoxon rank-sum test with Bonferroni correction.

Beyond macrophages, TIS also modulated adaptive lymphoid cells. Early memory B-cell clusters showed transcriptional activation of antigen presentation and proliferation (Figures S9C–S9E), and DCs mirrored proliferating macrophage signatures with enrichment in antigen-processing and metabolic-support pathways (Figures S9F–S9H). Although eosinophils were robustly recruited *in vivo*, single-cell analysis captured very few, likely due to technical loss or low transcript counts (reflected by low nCount per cell in Figure 3A). To overcome this limitation, bulk RNA-seq was performed on purified eosinophils from PBS- and TIS-treated mice (Figure 4A). Purity was validated by RT-qPCR, as sorted eosinophils expressed high levels of *Gata2*, *Ccr3*, *Siglecf*, *Epx*, and *Prg3* but lacked the T cell marker *Cd3* (Figure S10).

**Figure 4.**
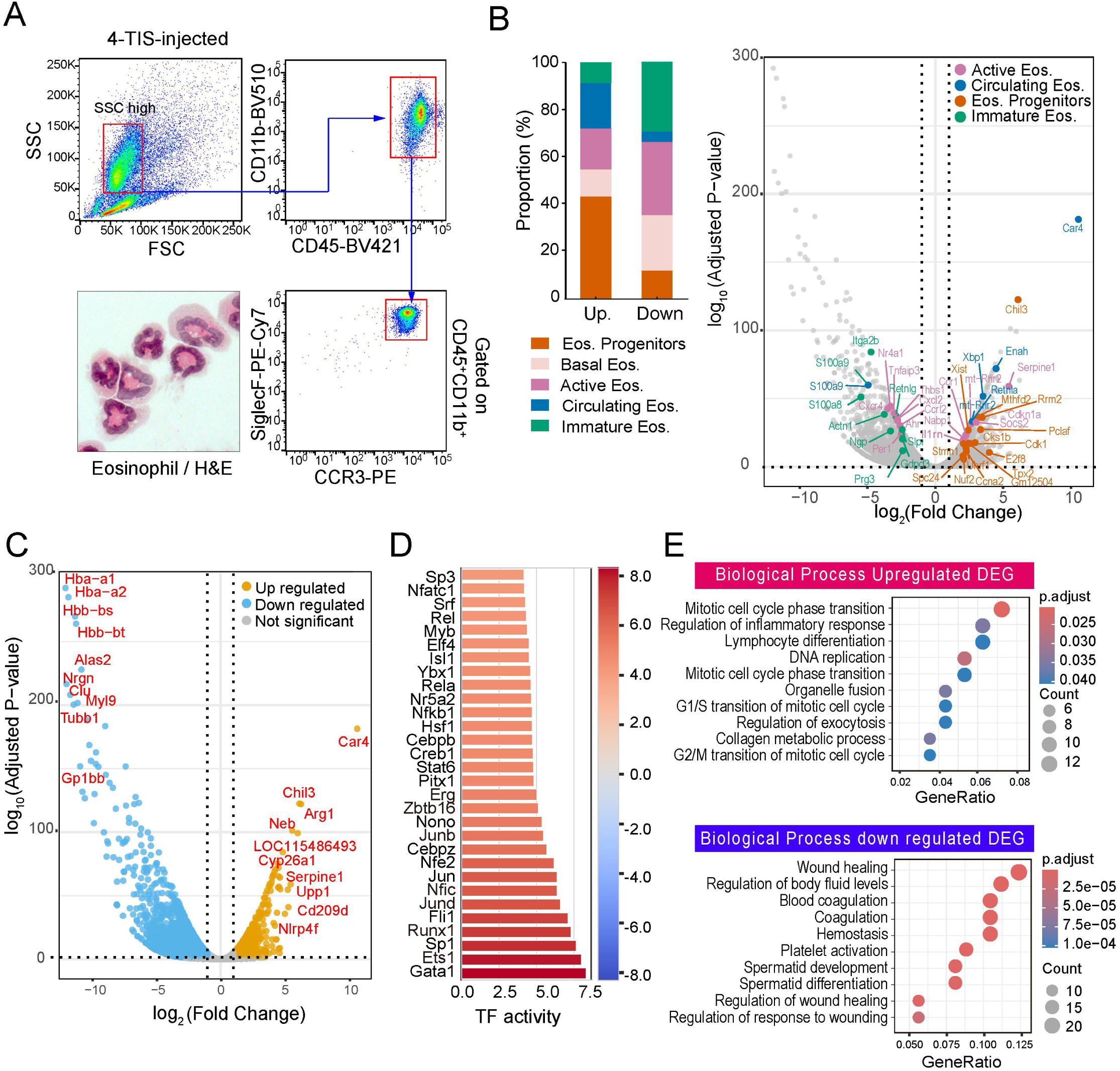
Bulk RNA-seq analysis shows that eosinophils adopt proliferative and immature-like transcriptional programs following 4-TIS treatment. (A) Flow cytometry gating strategy for eosinophil isolation from PBS- and 4-TIS–treated mice, with representative cytospin images (H&E). (B) Left: distribution of up- and down-regulated genes across eosinophil subtypes (progenitor, basal, circulating, immature). Right: volcano plot showing log₂ fold change vs – log₁₀ adjusted *P*. Genes with |log₂FC| ≥ 1 and adjusted *P* < 0.01 are highlighted; subtype markers are labeled. (C) Volcano plot of DEGs (4-TIS vs PBS) showing upregulated proliferative/metabolic genes (*Car4, Chil3, Arg1, Serpine1, Upp1*) and downregulated mature/erythroid genes (*Hba-a2, Hbb-bs, Alas2, Gp1bb*). (D) Inferred transcription factor activities enriched in DEGs, highlighting regulators of proliferation and lineage differentiation (*Runx1, Sp1, Ets1, Gata1*). (E) GO enrichment of DEGs. Top: biological processes upregulated by 4-TIS (e.g., mitotic cell cycle, DNA replication, lymphocyte differentiation). Bottom: downregulated processes (e.g., coagulation, platelet activation, wound healing). Data represent mean ± SD from two independent experiments; adjusted *P* values computed using DESeq2 with Benjamini– Hochberg correction.

Bulk RNA-seq revealed broad transcriptional remodeling of TIS-induced eosinophils ^19^. Among DEGs with ≥5-fold change, upregulated transcripts were enriched in progenitor/immature signatures, while downregulated genes were biased toward basal and active eosinophil markers (Figure 4B, left). Volcano plot annotation confirmed that proliferative genes such as *Car4*, *Chil3*, *Arg1*, *Serpine1*, *Upp1*, and *Ccna2* were predominantly linked to progenitor/immature subsets, while suppressed genes corresponded to mature effector states (Figure 4B, right). Across the full DEG set, upregulated transcripts included proliferative and metabolic regulators (*Car4*, *Chil3*, *Arg1*, *Serpine1*, *Upp1*), whereas downregulated genes were enriched for erythroid and hemostasis-related transcripts (*Hba-a1*, *Hbb-bs*, *Alas2*, *Gp1bb*) (Figure 4C). Transcription factor analysis revealed activation of regulators associated with proliferation and eosinophil lineage differentiation, including *Runx1*, *Sp1*, *Ets1*, and *Gata1* (Figure 4D). Gene ontology enrichment confirmed upregulation of mitotic cell cycle, DNA replication, and lymphocyte differentiation pathways, while genes related to tissue repair, coagulation, and hemostasis were downregulated (Figure 4E). Together, these findings indicate that TIS reprogram eosinophils toward a proliferative, metabolically active state while suppressing coagulation and tissue-repair functions. Collectively, TIS induce a delayed but broad immune response characterized by proliferating macrophages, activated B cells and DCs, and proliferative eosinophils, establishing a key axis of TIS-mediated immune modulation.

### 4-TIS suppresses MC38 tumor development and induces durable, cross-species anti-tumor immunity

To assess the anti-tumor efficacy of TIS, we employed a peritoneal MC38 colon carcinoma model. Mice received i.p. injections of TIS or LPS on days 0 and 7, were challenged with MC38 cells on day 9, and given a final dose on day 12 (Figure 5A). By day 23, bioluminescence imaging and gross examination revealed that TIS almost completely prevented tumor formation, whereas PBS- and LPS-treated animals developed large tumor burdens (Figure 5B). Tumor weights were markedly reduced in the TIS group. Peritoneal lavage analysis further showed that TIS expanded T cells, B cells, DCs, eosinophils, and both SPMs and LPMs, while maintaining relative subset distributions similar to wild-type mice, indicating robust immune activation without disrupting homeostasis (Figures 5C and S11).

**Figure 5.**
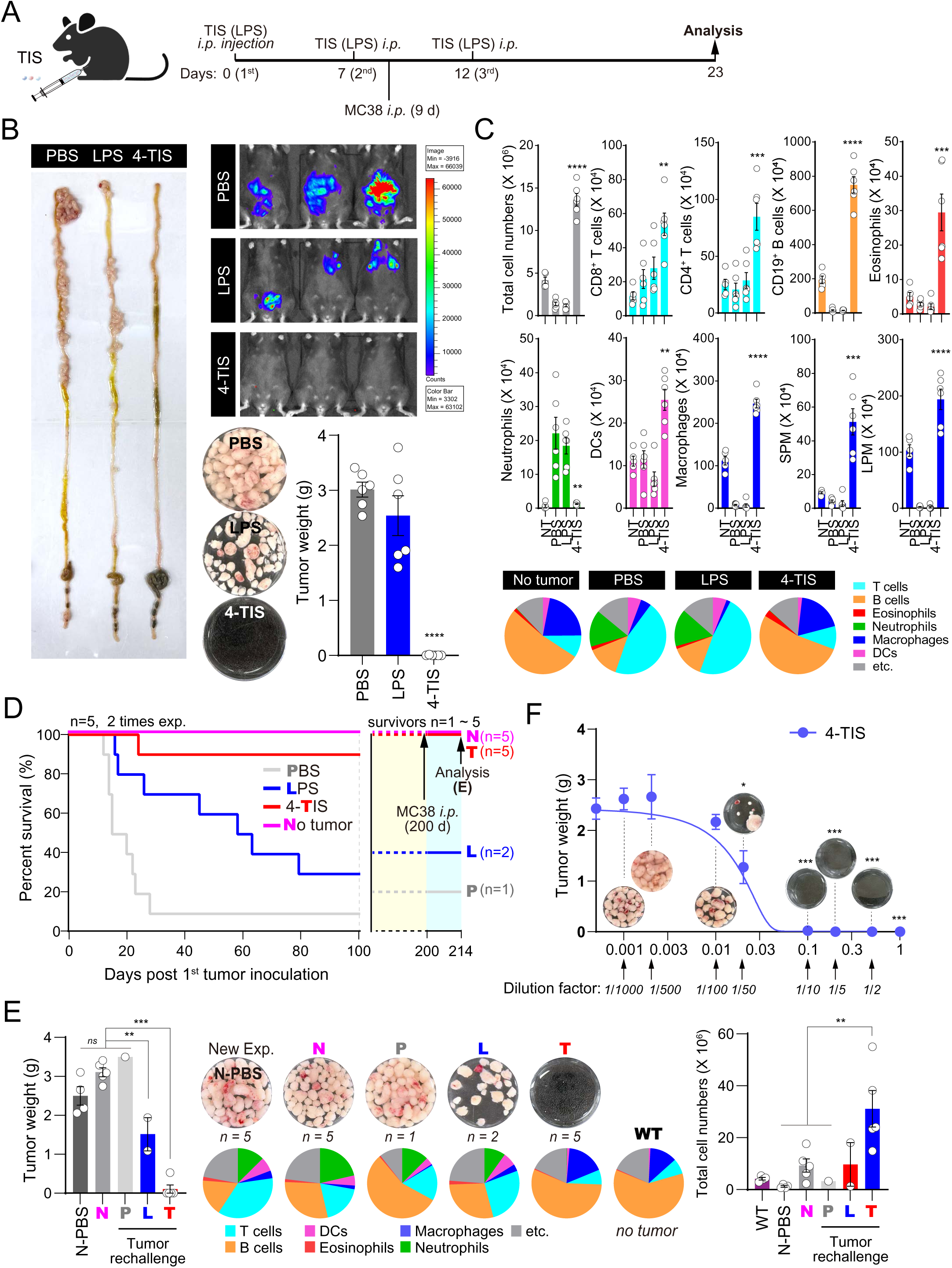
4-TIS suppresses MC38 colon cancer development *in vivo*. (A) Experimental design: i.p. injection of PBS, LPS (25 µg), or 4-TIS on days 0, 7, and 12, followed by MC38 tumor inoculation on day 9 and analysis on day 23. (B) Tumor burden at day 14. Left: representative intestinal tracts. Middle: bioluminescence imaging. Right: representative excised tumors and weights (mean ± SEM, ****p < 0.0001 vs. PBS). (C) Peritoneal immune composition. Total leukocytes and subsets (CD8⁺ T, CD4⁺ T, CD19⁺ B, DCs, eosinophils, neutrophils, macrophages, SPMs, LPMs). Pie charts depict CD45⁺ cell proportions. (D) Kaplan–Meier survival. Mice (n = 5 per group) were monitored for 200 d. Surviving mice were rechallenged with MC38 at day 200 (arrows). (E) Outcomes after rechallenge. Tumor images, weights, total cell counts, and immune distributions in survivors (N: no tumor; P, L, T: survivors of PBS, LPS, 4-TIS). (F) Dose–response of TIS. Serial dilutions of 4-TIS were tested; endpoint tumor weights and representative nodules shown. Data are mean ± SEM; *p < 0.05, **p < 0.01, ***p < 0.001, ****p < 0.0001.

TIS also conferred durable protection. In survival studies, TIS-treated mice remained tumor-free for >100 days, with significant survival advantage over PBS- and LPS-treated controls (Figure 5D). Long-term monitoring up to 200 days showed that all surviving TIS-treated mice resisted MC38 rechallenge without additional treatment, whereas controls exhibited rapid tumor outgrowth (Figures 5D and 5E). Endpoint analysis revealed almost complete absence of peritoneal tumors in TIS survivors, in contrast to extensive nodules in control groups. Flow cytometry confirmed that TIS survivors maintained balanced expansion of T cells, B cells, DCs, eosinophils, and macrophages, consistent with durable and functional anti-tumor immunity (Figure 5E). Tumor suppression correlated with TIS dose, with serial dilutions producing a clear dose-dependent reduction in tumor burden (Figure 5F).

The persistence of protection was further validated in staggered booster experiments mimicking vaccination schedules (Figure 6A). Mice receiving two priming doses plus a third booster at 1 or 3 months remained completely tumor-free after MC38 challenge. Even in the 4-month cohort that received only two doses, protection was maintained (Figure 6B). In contrast, PBS- and LPS-treated animals consistently developed tumors. Endpoint analyses revealed expansion of multiple immune subsets without skewing overall composition (Figures S12B–S12E), indicating that TIS reinforces peritoneal immunity while preserving balance.

**Figure 6.**
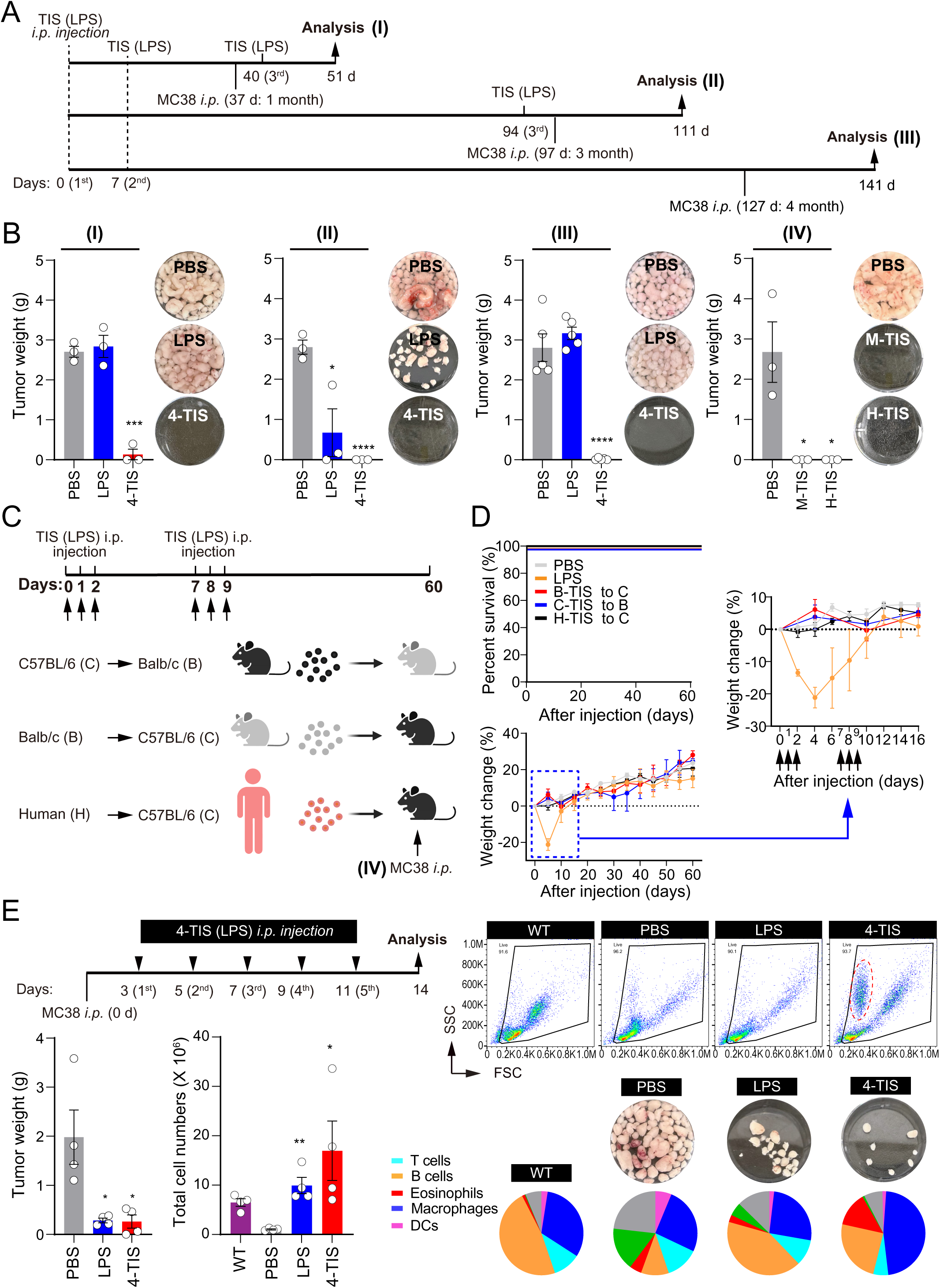
4-TIS confers long-term preventive and therapeutic tumor protection. (A) Schedules of delayed challenge. Mice received 4-TIS on days 0 and 7, followed by: (I) MC38 on day 37 with a third TIS on day 40; (II) third TIS on day 94 then MC38 on day 97; (III) MC38 on day 127 without a third dose. (B) Tumor weights and representative images at day 14 post-challenge. Panels I–III: schedules in (A). Panel IV: xenogeneic transfer of human-derived TIS (H-TIS) into C57BL/6 mice. (C) Cross-species safety. Experimental design: C57BL/6↔Balb/c allogeneic TIS, or human→C57BL/6 xenogeneic TIS. (D) Survival and weight. Allogeneic and xenogeneic TIS caused no adverse effects, unlike LPS. Mice receiving H-TIS remained protected after MC38 challenge. (E) Therapeutic efficacy. Mice inoculated with MC38 (day 0) received 4-TIS on days 3–11. At day 14, tumor burden, total peritoneal cell counts, flow cytometry (FSC/SSC), and pie charts of immune subsets are shown. Red dashed box highlights SSChigh eosinophils. Bars show mean ± SEM; *p < 0.05, **p < 0.01, ***p < 0.001, ****p < 0.0001.

TIS also demonstrated excellent safety and cross-species efficacy. Allogeneic TIS administration caused no weight loss or mortality, whereas LPS induced ≥20% body weight loss (Figures 6C and 6D). Xenogeneic human TIS injected into C57BL/6 mice caused no adverse effects and, remarkably, provided complete protection against subsequent MC38 challenge, while PBS-treated controls developed tumors (Figure 6B, group IV). Thus, human-derived TIS are not only safe but also confer durable cross-species protection. Therapeutic administration of TIS after tumor inoculation significantly reduced tumor mass by day 14, demonstrating efficacy even against established disease (Figure 6E). Collectively, these findings demonstrate that TIS potently suppress MC38 tumors, induce rechallenge-resistant immunity, expand immune subsets without disrupting homeostasis, remain safe in allogeneic and xenogeneic contexts, and retain therapeutic activity against established disease.

Beyond the peritoneal setting, we asked whether TIS could provide systemic protection against distant metastasis. Intradermal TIS administration before i.v. B16F10 melanoma challenge markedly reduced pulmonary metastases compared with PBS, as measured by lung nodules and histology (Figures S13A–S13C). Whereas LPS induced local inflammation and tissue damage, TIS caused no visible lesions, highlighting its safety (Figure S13B). Systemic protection correlated with increased splenic T and B cells, expansion of CXCR5⁺ Tfh-like and CD44⁺ memory T cells, and elevated serum IL-2 and IFN-γ (Figures S13D and S13E). These results show that TIS-mediated tumor suppression extends beyond the peritoneal cavity to distant metastatic sites.

### Macrophages and eosinophils are critical mediators of 4-TIS-induced tumor suppression

Given the pronounced remodeling of peritoneal macrophages and eosinophils after TIS administration, we next asked whether these cell types are essential for anti-tumor efficacy. Macrophage depletion with clodronate liposomes markedly impaired tumor control when only two TIS doses were given, leading to increased tumor burden, reduced leukocyte accumulation, and altered immune composition (Figures 7A–7C). Notably, however, the addition of a third TIS dose restored anti-tumor activity even under macrophage depletion (Figures 7D and 7E), indicating that newly recruited monocytes replenish the macrophage pool and can be reprogrammed by TIS to sustain tumor suppression.

**Figure 7.**
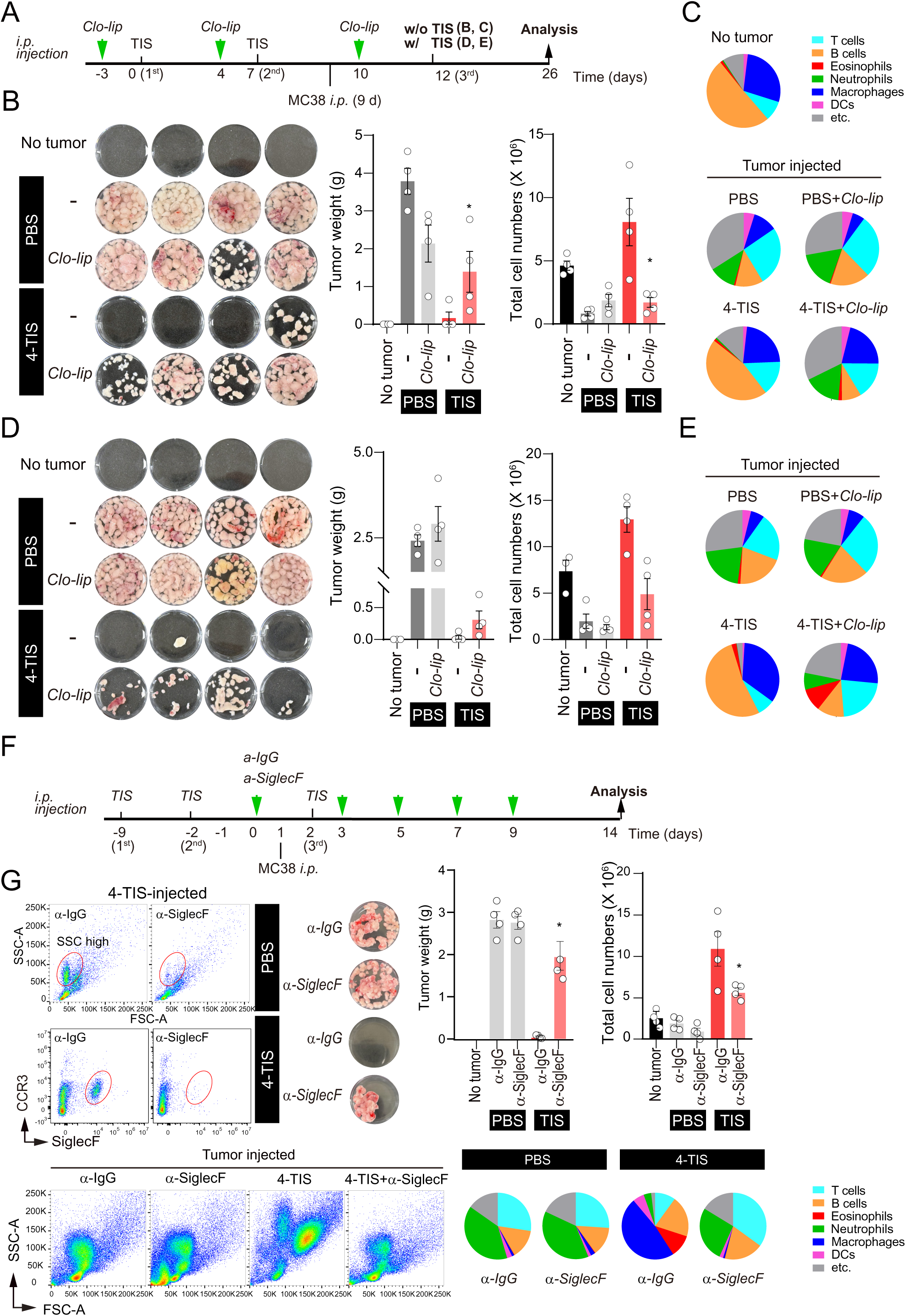
Macrophage and eosinophil depletion diminishes 4-TIS antitumor efficacy. (A) Experimental design for macrophage depletion (clodronate liposomes, Clo-lip). (B–C) Without a third TIS dose: excised tumors, weights, total peritoneal leukocytes, and CD45⁺ immune composition. Clo-lip abrogated tumor suppression. (D–E) With a third TIS dose: tumor control and immune composition largely restored, suggesting recruitment and reprogramming of new macrophages. (F) Experimental design for eosinophil depletion (α–Siglec-F antibody). (G) Flow cytometry confirmed eosinophil loss. Tumor weights, total leukocytes, and immune composition at day 14 show that eosinophil depletion almost completely abolished TIS-mediated protection. Data are mean ± SEM. *p < 0.05, **p < 0.01, ***p < 0.001, ****p < 0.0001.

In case of eosinophil depletion with anti-Siglec-F antibodies, TIS-mediated protection was almost completely lost, resulting in dramatic tumor progression, failure of leukocyte expansion, and profound disruption of immune balance (Figure 7G). Flow cytometric analyses confirmed that eosinophils are indispensable for TIS-induced tumor control, in sharp contrast to the conditional requirement of macrophages. Together, these findings establish that TIS act as a powerful cell-free immune modulator that induces durable tumor immunity through a macrophage–eosinophil axis, thereby linking T cell–derived vesicle biology to functional anti-tumor mechanisms.

## DISCUSSION

Innate immune responses are characterized by their immediate and often excessive reactions to a wide variety of stimuli. This rapid response is essential for eliminating invading microbes or viruses during the earliest phase of infection. Those that escape this initial defense are left to be selectively and precisely targeted by the adaptive immune system ^21^. Given that adaptive immunity operates with high specificity toward particular antigens, it is unlikely to generate the kind of forceful, systemic response commonly observed in innate immunity. Instead, its actions are more focused and regulated. In this study, our fundamental question was: What happens when large quantities of T cell–originated materials—the core effectors of adaptive immunity—are introduced into the body in the absence of specific antigens? Because this approach bypasses conventional antigen recognition, its potential effects remain unpredictable and have never been explored before.

Our findings challenge the conventional view of adaptive immunity by introducing TIS—cell-free vesicles derived from T-cell microvilli during immune synapse formation—as a novel modality that mimics and amplifies adaptive immune signals without direct T-cell contact. TIS carry a naturally packaged set of T cell–derived membrane proteins, signaling molecules, and metabolic cues that are efficiently phagocytosed by macrophages and DCs. Their uptake reprogrammed these innate cells toward a non-inflammatory, adaptive-like phenotype, in contrast to the acute inflammatory cascades elicited by canonical innate stimuli such as LPS. Functionally, this resulted in delayed but sustained effector recruitment, potent tumor suppression, and durable immune activation lasting for months. These findings suggest that TIS act as active messengers between innate and adaptive immunity, enabling memory-like responses without classical antigen presentation or systemic inflammation.

Single-cell and bulk transcriptomic profiling revealed that TIS reprogram Timd4⁺ large peritoneal macrophages into metabolically active and proliferative phenotypes while suppressing tolerogenic programs. Compared to PBS or LPS, TIS upregulated the C/EBP family (*Cebpa, Cebpb, Cebpg, Cebpd*) and *Ncoa1*, genes associated with metabolic activation and partial M1-like polarization ^22,23^, while repressing regulatory and tissue-resident factors such as *Irf8, Myb,* and *Gata3* ^24,25^. Gene signatures further highlighted antigen presentation (*Clec4d, C1qc*), cellular metabolism (*Ucp1, Gst*), and translational machinery (*Rps24*), collectively indicating a shift from a homeostatic to an immune-stimulatory state. Network analyses pointed to two dominant modules: metabolic and antioxidant programs (*Atf6, Ppargc1a, Mef2a, Nr3c1*) ^26^, and anti-inflammatory/tissue-repair pathways (*Il6, Il1rn, Acot13, C3*) ^27^, suggesting that TIS prime macrophages for sustained activation while maintaining tissue balance. A striking feature of TIS-treated peritoneum was the emergence of proliferating macrophages, absent in PBS or LPS conditions, marked by induction of cell-cycle and DNA replication genes (*Mki67, Top2a, Birc5, Cenpf, Ccnb2*) ^28^. This proliferative response, together with eosinophil remodeling, underpins the delayed but broad accumulation of myeloid and adaptive immune populations observed in vivo and provides a mechanistic basis for the durable tumor suppression elicited by TIS.

These cellular programs not only explain the immediate antitumor activity of TIS but also suggest that they imprint longer-term memory-like properties in innate cells. Indeed, a particularly intriguing observation is that TIS confer protective immunity for over six months—despite no co-administered antigen—implying that innate rather than classical adaptive memory mechanisms are at play. Consistent with this, our transcriptomic analyses revealed upregulation of *Ptgs2* (COX-2), implicating metabolic and epigenetic remodeling characteristic of trained immunity ^29^. Concurrent induction of IFN-γ–responsive GTPases (*Gbp2b, Gbp4, Gbp11*) reinforces this notion, as GBPs are key effectors in innate immune memory and pathogen sensing ^30^. Crucially, transcription factors such as *Cebpb*, which are required for trained immunity in hematopoietic stem cells through chromatin remodeling, were also enriched in TIS-treated macrophages ^31^. Together with the proliferative and metabolic programs observed in eosinophils, these molecular signatures support a model wherein TIS instigate a durable, trained state in innate cells. This mechanism likely underlies the prolonged anti-tumor efficacy of TIS without repeated antigen exposure and highlights their potential as an immunotherapeutic platform that leverages innate immune memory for sustained protection ^34^.

A key therapeutic advantage of TIS lies in their intrinsic targeting to professional APCs, primarily macrophages and DCs. These are the very cells that T cells naturally engage to form IS in vivo, meaning that TIS inherently deliver T cell–derived signals to the correct cellular recipients without requiring any artificial genetic modification or engineered targeting system. This property makes TIS an attractive platform for immunotherapy. Moreover, because immunological synaptosomes are shed predominantly from T cell microvilli, engineering the expression of selected cytokines or ligands on microvilli could further enhance TIS functionality. When displayed on the TIS membrane, these molecules would benefit from avidity effects, potentially achieving far greater bioactivity than their soluble monomeric counterparts. Another notable feature of TIS is their remarkable persistence *in vivo*. Unlike peptides, small molecules, or soluble cytokines—which typically have half-lives ranging from minutes to a few hours—TIS remain in the body for extended periods, allowing sustained immunological effects. Efficiently phagocytosed by macrophages and DCs, TIS enable stable delivery of T cell–derived signals, leading to sustained myeloid activation and prolonged cytokine release that fosters durable adaptive immune responses.

Recent studies have highlighted the antitumor potential of T cell–derived EVs, including mechanically generated nanovesicles and CAR-T cell–derived exosomes ^32,33,35^. These strategies, however, have primarily emphasized the delivery of cytotoxic molecules or the blockade of immunosuppressive pathways such as PD-1/PD-L1 and TGF-β. They are also generally limited to single-dose or short-term evaluations and often depend on antigen specificity, restricting their capacity to capture the broader dynamics of adaptive immunity. In addition, most studies have relied on EVs collected from culture supernatants ^32,33,35^, whereas our work focuses on T-cell immunological synaptosomes (TIS)—vesicles generated within an hour of immune synapse formation and strictly dependent on adhesion. By administering TIS sequentially in vivo, we uncovered functional properties that were not evident in short-term or single-injection studies. TIS orchestrated a coordinated cellular network involving macrophages, eosinophils, and lymphocytes, leading to the establishment of trained immunity–like programs and durable systemic tumor suppression.

In conclusion, this study demonstrates the potential of TIS as a novel immunomodulatory platform capable of inducing durable anti-tumor responses with minimal toxicity. While this work focused on CD4⁺ T cell–derived TIS, future studies should evaluate TIS from CD8⁺ T cells, which may offer additional cytotoxic or regulatory functions. Broader assessment across cancer types and administration routes will be essential to optimize clinical translation. Importantly, TIS drive long-term immune reprogramming without disrupting host homeostasis, suggesting suitability for sustained use. Their defined cellular origin offers opportunities for therapeutic enhancement—for example, by engineering donor T cells to express specific cytokines or antigens on microvilli, enabling precise immune modulation. With appropriate dosing and immunogenetic compatibility, TIS could serve as an “off-the-shelf” cancer immunotherapy for both treatment and recurrence prevention.

## Material and Method

### Reagents and antibodies

For flow cytometry, Fcγ receptor–blocking antibody (clone 2.4G2; Cat# 553141) was obtained from BD Biosciences (San Jose, CA, USA). Fluorochrome-conjugated antibodies against Ly6G, CD11b, CD3ε, CD8, CD4, CD45, CD19, CD11c, MHC class II (I-A/I-E), F4/80, Siglec-F, CCR3, CD40, CD80, and CD86 were purchased from eBioscience (San Diego, CA, USA) and Biolegend (San Diego, CA, USA). All flow cytometry antibodies were used at a dilution of 1:200. For Western blotting, primary antibodies against ARRDC1 (Abcam, Cambridge, UK) and TCRζ (Santa Cruz Biotechnology, Dallas, TX, USA) were used at a dilution of 1:1000. Lipopolysaccharide (LPS) was obtained from Sigma-Aldrich (St. Louis, MO, USA). Clodronate liposomes were purchased from Liposoma (Amsterdam, Netherlands). Anti-CD3/CD28 antibody–conjugated CNBr-activated Sepharose beads were prepared as previously described ^18^. Recombinant murine ICAM-1/CD54 was purchased from R&D Systems (Minneapolis, MN, USA). MHC class II (I-Ab) monomers were provided by the NIH Tetramer Core Facility (Emory University, Atlanta, GA, USA). PMS-green cell mask and CellTrace™ Violet (CTV) Cell Proliferation Kit were purchased from Thermo Fisher Scientific (Waltham, MA, USA). Mouse TNF-α DuoSet ELISA kits, anti– Siglec-F monoclonal antibody, and rat anti-IgG2A Isotype control antibody were purchased from R&D Systems (Minneapolis, MN, USA). Anti-mouse CD3, anti-mouse CD28, anti-human CD3, anti-human CD28, anti-TNFαR, and anti-GM-CSFR antibodies were purchased from Bio X Cell (Lebanon, NH, USA). D-Luciferin (potassium salt) was purchased from PerkinElmer (Waltham, MA, USA). MojoSort™ Mouse Pan B Cell, CD4⁺ T Cell, and CD8⁺ T Cell isolation kits, as well as LEGENDplex™ cytokine/chemokine kits, were purchased from BioLegend (San Diego, CA, USA). Polyinosinic-polycytidylic acid [Poly(I:C)] was purchased from InvivoGen (San Diego, CA, USA).

### Cells

MC38 colon carcinoma cell lines (YC-A002, Ubigene) and luciferase-expressing MC38 cells (MC38-luc, YC-A002-Luc-P, Ubigene) were cultured in RPMI-1640 medium (Invitrogen, Carlsbad, CA, USA) supplemented with 10% fetal bovine serum (FBS; Invitrogen) and 1% penicillin–streptomycin (Gibco, Waltham, MA, USA). Naïve CD3⁺, CD4⁺, and CD8⁺ T cells were isolated from spleens and lymph nodes of C57BL/6 or OT-II TCR transgenic mice using MojoSort™ negative selection magnetic cell separation kits (BioLegend), according to the manufacturer’s protocol. Briefly, single-cell suspensions were obtained by mechanical dissociation of spleen and lymph node tissues, followed by filtration through 40 µm strainers (Corning, NY, USA), centrifugation (2000 × g, 5 min), and red blood cell lysis (RBC Lysis Buffer; eBioscience). To generate mouse T-cell blasts, purified CD3⁺, CD4⁺, or CD8⁺ T cells from C57BL/6 mice were stimulated for 48 h on culture plates coated with 2 µg/mL anti-CD3/CD28 antibodies in the presence of recombinant IL-2 (100 U/mL), followed by continued culture for an additional 5 days with rIL-2 to promote expansion. B-cell blasts were generated by stimulating CD19⁺ B cells from C57BL/6 spleens with LPS (10 µg/mL) for 3 days in complete RPMI-1640 medium. GM-CSF–differentiated bone marrow cells (BMCs) were generated by flushing bone marrow from femurs and tibias of C57BL/6 mice, followed by culture in complete RPMI supplemented with 20 ng/mL recombinant murine GM-CSF (Sino Biological, Beijing, China). Fresh medium was added on days 3 and 5, and cells were harvested on day 7 for downstream assays. In some experiments, CD11c⁺ DCs or F4/80⁺ macrophages derived from bone marrow cultures were isolated using magnetic separation kits (Miltenyi Biotec, Bergisch Gladbach, Germany). Peritoneal macrophages were harvested 3 days after thioglycollate injection. Human CD3⁺ T cells were purchased from Lonza (Basel, Switzerland) and used for TIS preparation.

### Animals

C57BL/6 wild-type mice, OT-II TCR transgenic mice (C57BL/6 background), and BALB/c mice were obtained from The Jackson Laboratory (Bar Harbor, ME, USA) and maintained under specific pathogen-free (SPF) conditions. All animal experiments were conducted in accordance with protocols approved by the Institutional Animal Care and Use Committee of the School of Life Sciences, Gwangju Institute of Science and Technology (Gwangju, Korea) and carried out in accordance with their approved guidelines (IACUC GIST-2023-048, GIST-2024-022, GIST-2023-001).

### TIS isolation and lyophilization

TIS were isolated following a modified protocol based on previously published methods ^18^. Briefly, 5 × 10^7^ CD4^+^ T blasts from C57BL/6 wild-type mice were resuspended in 5 mL of RPMI 1640 medium supplemented with 10% exosome-depleted fetal bovine serum (exo-free FBS; Gibco), and incubated with 0.5 mL of a 50% slurry of anti-CD3/CD28 antibody-coated sepharose 4B beads for 3 h at 37 °C in a CO₂ incubator. After incubation, the bead–cell mixture was vigorously vortexed for 1 min and centrifuged. Supernatants were sequentially centrifuged at 2000 × g for 10 min and 2500 × g for 10 min to remove residual cells. The clarified supernatant was then ultracentrifuged at 100,000 × g for 1 h at 4 °C, and the resulting TIS-containing pellet was resuspended in sterile PBS and stored at –70 °C until use. To isolate TIS from human T cells, CD3^+^ T blasts were incubated with anti-CD3/CD28– coated beads as described above. For long-term storage and reconstitution, freshly isolated TIS were subjected to lyophilization. Samples were flash-frozen in liquid nitrogen and lyophilized overnight under vacuum at –50 °C using a laboratory freeze-dryer (Operon). Lyophilized TIS (L-TIS) were reconstituted in sterile PBS immediately before use.

### Quantitation of TIS

To assess TIS release under different stimulation conditions, PMS green–labeled T cells (5 × 10⁵ cells per well) were suspended in L-15 medium and seeded into 12-well plates. Cells were stimulated under one of four conditions: control (uncoated), bead-bound anti-CD3/CD28, peptide–MHC/ICAM-1–coated, or soluble anti-CD3/CD28 in uncoated wells. At 0, 10, 30, 60, and 180 min post-stimulation, culture supernatants were collected and sequentially centrifuged at 5000 rpm for 3 min twice, followed by a final spin at 8000 rpm for 3 min to remove residual cells and debris. Supernatants containing TIS were analyzed on a CytoFLEX flow cytometer ((Beckman Coulter, Brea, CA, USA) configured for small particle detection. The 405/10 nm violet side scatter (VSSC) filter was reassigned to the V450 channel in the wavelength division multiplexer, and detector settings were adjusted accordingly in CytExpert software (Beckman Coulter). The event detection threshold was set on the VSSC-Height channel and manually optimized using the Flow Cytometry Submicron Particle Size Reference Kit (Thermo Fisher Scientific) at the recommended dilution to ensure accurate detection of submicron vesicles.

### In vitro stimulation of BMDCs, B cells, and macrophages with TIS

Each cell type was seeded into appropriate culture plates and stimulated for 24 h with PBS, LPS (100 ng/mL for BMDCs and macrophages; 1 μg/mL for B cells), or TIS derived from CD3⁺, CD4⁺, or CD8⁺ T cells in fresh, frozen, or lyophilized form. For all in vitro assays, TIS were applied at a 1:40 ratio relative to target cell numbers. After stimulation, BMDCs and B cells were stained with antibodies against CD11c or CD19 and MHC class II and analyzed by flow cytometry. For macrophages, TNF-α levels in culture supernatants were quantified by ELISA, and nitrite production was measured using the Griess assay.

### LEGENDplex bead-based immunoassay

Skin and plasma levels of indicated cytokines or chemokines were quantified using LEGENDplex mouse proinflammatory cytokine or chemokine (13-plex, 8-plex) according to the manufacturer’s instructions.

### Western blot analysis

To confirm the molecular composition and cellular origin of isolated TIS, Western blotting was performed to detect ARRDC1 and TCRζ. TIS were isolated and fractionated on a sucrose gradient (0.2–2.0 M) as previously described. Equal amounts of protein from pooled fractions were separated by SDS–PAGE and transferred to nitrocellulose membranes (Millipore, Billerica, MA, USA). Membranes were probed with primary antibodies against ARRDC1 and TCRζ, followed by HRP-conjugated secondary antibodies. Signals were visualized using enhanced chemiluminescence (ECL) reagents (Amersham, GE Healthcare, Chicago, IL, USA).

### Microarray-Based Gene Expression Analysis

Day 9 BMCs were treated with PBS, LPS, TIS, or LPS+TIS for 24 h. Total RNA was extracted using the RNeasy Mini Kit (Qiagen), and RNA quality was assessed with a Bioanalyzer 2100 (Agilent Technologies). Gene expression profiling was performed using the GeneChip™ Mouse Gene 2.0 ST Array (Affymetrix, Thermo Fisher Scientific). cDNA synthesis, fragmentation, and biotin labeling were performed according to the manufacturer’s protocol. Arrays were hybridized at 45 °C for 16 h, washed, stained, and scanned using the Affymetrix system. Raw CEL files were processed with Affymetrix Power Tools (APT), and expression data were normalized by the robust multi-array average (RMA) method. Control and non-informative probes were filtered, leaving 33,793 probes for analysis. Differential gene expression was determined based on fold change. Hierarchical clustering was performed using Euclidean distance with complete linkage. Gene ontology and pathway enrichment analyses were conducted using g:Profiler and KEGG databases. All analyses and data visualization were performed in R.

### Analysis of peritoneal immune cell dynamics following TIS administration

To examine the effects of TIS on peritoneal immune cell composition, C57BL/6 mice were injected i.p. with PBS, LPS (25 μg/mouse), poly(I:C) (50 μg/mouse), or TIS on days 0, 7, and 10. Peritoneal cells were collected by lavage at 4 h, 1 day, 2 days, 8 days, and 11 days after the initial injection and analyzed by flow cytometry. The experimental timeline is illustrated in Figure 2A. Gating was performed on live, singlet CD45⁺ cells, and subsequent analysis quantified T cells, B cells, macrophages, DCs, neutrophils, and eosinophils over time. For all in vivo assays, each mouse received a single intraperitoneal dose equivalent to the yield from 5 × 10⁷ T cells.

### Analysis of skin immune cell recruitment following TIS administration

To evaluate immune cell recruitment in the skin, C57BL/6 mice were injected intradermally (i.d.) with PBS, LPS (25 μg/mouse), poly(I:C) (50 μg/mouse), or TIS on days 0, 7, and 10. Skin tissues at the injection sites were harvested at 1, 2, 8, and 11 days after the initial injection. The experimental timeline is illustrated in Figure S6A. Samples were fixed in 10% neutral-buffered formalin, embedded in paraffin, sectioned, and stained with hematoxylin and eosin (H&E). Infiltrating immune cells, including macrophages, lymphocytes, neutrophils, and eosinophils, were identified by histological criteria and quantified in representative fields (cells/high-power field). Skin thickness was measured using digital microscopy. Chemokine levels (CCL24, CCL9, CCL7) in skin homogenates and serum were quantified using LEGENDplex™ Mouse Cytokine/Chemokine Panel according to the manufacturer’s instructions.

### In vivo uptake analysis of TIS by peritoneal immune cells

To evaluate the kinetics of TIS uptake in vivo, PMS Green–labeled TIS were injected i.p. into C57BL/6 mice. Peritoneal cells were collected by lavage at 0.5, 4, 6, 24, 48, and 72 h post-injection and analyzed by flow cytometry. Gating was performed on CD45⁺ cells to determine uptake across peritoneal leukocyte subsets, including macrophages (F4/80⁺CD11b⁺), DCs (CD11c⁺MHCII⁺), B cells (CD19⁺), T cells (CD3⁺), and eosinophils (Siglec-F⁺). To assess the impact of repeated dosing, C57BL/6 mice were injected i.p. with unlabeled TIS on days 0 and 7, followed by PMS Green–labeled TIS on day 10 (third injection). 4 h later, peritoneal cells were collected by lavage and analyzed by flow cytometry. The frequency of TIS-positive populations was determined across macrophages, DCs, B cells, T cells, and eosinophils to evaluate whether prior exposure altered uptake dynamics.

### Visualization of TIS uptake by peritoneal macrophages

To visualize TIS uptake, peritoneal macrophages were isolated from thioglycollate-treated mice and labeled with CellTrace™ Violet (CTV) according to the manufacturer’s protocol. PMS Deep Red–labeled TIS were added to macrophage cultures and incubated at 37 °C. At the indicated time points, cells were imaged using an LSM880 confocal microscope equipped with an Airyscan super-resolution module (Zeiss). Image processing and colocalization analysis of TIS signals with intracellular structures were performed using FluoView software (Olympus).

### Realtime quantitative PCR

Peritoneal cells were collected 24 h after the third i.p. TIS dose by lavage (5–8 mL PBS, 2% FBS). Cells were plated in 6-well plates at 2× 10⁶ cells/well in RPMI-1640 + 10% FBS and incubated 2 h at 37 °C, 5% CO₂ to enrich adherent macrophages. For non-adherent cells were removed twice PBS washes, and adherent cells were lysed. Eosinophils were isolated from the peritoneal lavage of C57BL/6 mice following three sequential intraperitoneal injections of TIS. Cells were enriched by flow cytometric sorting based on CD45⁺CD11b⁺Siglec-F⁺CCR3⁺ expression, whereas control T cells were obtained by sorting CD4⁺ splenocytes from naïve mice. Total RNA was extracted using the RNeasy Mini Kit (Qiagen, Hilden, Germany) and reverse-transcribed into cDNA using TOPscript™ RT DryMIX (Enzynomics) according to the manufacturer’s protocol. Real-time quantitative PCR (qPCR) was performed with LightCycler® 480 SYBR Green I Master Mix (Roche, Basel, Switzerland) on a LightCycler® 480 Real-Time PCR System (Roche). Primer sequences for *Cd3, Gata2, Ccr3, Siglec-f, Epx, Prg3, Ccl24, Ccna2, Mki67, Ccl4, Ifna1, and Itgb7* were obtained from the Harvard PrimerBank database and synthesized by Macrogen (Seoul, Korea). Primer sequences are provided in Table S1.

### Seahorse metabolic flux analysis

Sensor cartridges (96-well) were hydrated overnight at 37 °C in a non-CO₂ incubator with 200 µl of XF Calibrant solution (Agilent Technologies, Santa Clara, CA, USA). XF RPMI assay medium (Agilent) was supplemented with 10 mM glucose, 1 mM pyruvate, and 2 mM L-glutamine. Seahorse XFp culture plates (Agilent) were coated with 0.01% poly-L-lysine (PLL; Sigma-Aldrich, St. Louis, MO, USA) overnight at 4 °C, and PLL was removed immediately before cell seeding. For macrophage assays, peritoneal cells (5 × 10⁵/well) were seeded onto PLL-coated plates, incubated for 2 h at 37 °C, washed twice with PBS, and enriched for adherent macrophages. For B cell assays, peritoneal B cells were isolated using the MojoSort™ Mouse Pan B Cell Isolation Kit, stabilized for 1 h at 37 °C, and seeded at the same density. Basal oxygen consumption and acidification rates were recorded for 24 min, followed by sequential injections of metabolic inhibitors (final volume, 200 µl) oligomycin (1.5 µM), FCCP (2 µM), and rotenone/antimycin A (0.5 µM). Stimulated rates were monitored for an additional 2 h.

### Single-cell RNA sequencing of peritoneal immune cells

Peritoneal immune cells were isolated from C57BL/6 mice treated with PBS, LPS, or 4-TIS. Cells were washed and resuspended in 0.04% BSA/DPBS, with an average viability >85% prior to library construction. Approximately 25,000 cells per sample were loaded into the Chromium Controller (10x Genomics), targeting ∼16,000 recovered cells per library. Libraries were generated using the Chromium GEM-X Single Cell 3′ Kit v4 according to the manufacturer’s instructions. cDNA was amplified (11 cycles), followed by sample index PCR (14 cycles) using sample-specific barcodes (PBS: TT-E4; LPS: TT-E5; 4-TIS: TT-E7). Final libraries yielded ∼800–1,000 ng with concentrations ranging from 70–87 nM. Library size distribution and quality were assessed using a Bioanalyzer 2100 (Agilent Technologies), confirming optimal profiles for both cDNA and final libraries. Sequencing was performed on an Illumina short-read platform, generating paired-end reads (101 bp).

### Bulk RNA sequencing of TIS-induced eosinophils

Eosinophils were isolated from peritoneal lavage fluid of C57BL/6 mice that received three intraperitoneal injections of TIS (days 0, 7, and 10). PBS-treated mice served as controls. Eosinophils were identified by SSC^high^, CD45⁺CD11b⁺Siglec-F⁺CCR3⁺ expression, and confirmed by cytospin preparations followed by H&E staining. Purity was further validated by real-time qPCR for eosinophil marker genes (Gata2, Ccr3, Siglec-F, Epx, and Prg3). Total RNA was extracted using the RNeasy Mini Kit (Qiagen) and quality was assessed with a Bioanalyzer 2100 (Agilent Technologies). Library preparation and sequencing were performed by Macrogen (Seoul, Korea) using the SMART-Seq v4 Ultra Low Input RNA Kit and Nextera XT DNA Library Preparation Kit. Libraries were sequenced on the Illumina platform to generate 101-bp paired-end reads. Reads were aligned to the mouse reference genome (mm10) with NCBI_108 annotation. Downstream differential gene expression and pathway enrichment analyses were performed in R.

### In vivo tumor prevention and long-term protection assays

To evaluate the cancer-preventive effect of TIS, C57BL/6 mice received intraperitoneal injections of TIS on days 0, 7, and 12. On day 9, mice were challenged i.p. with MC38 colon carcinoma cells (3 × 10⁵ cells/mouse). On day 23, mice were euthanized, and peritoneal cells were collected by lavage for flow cytometric analysis of immune cell composition. Visible tumor nodules adhering to the peritoneal lining and intestinal surfaces were excised, weighed, and documented. For bioluminescence-based tumor monitoring, MC38-luc cells stably expressing firefly luciferase were used. Mice received an intraperitoneal injection of D-luciferin (3 mg/mouse), and whole-body luminescence was imaged using the IVIS Spectrum system (PerkinElmer). To assess dose dependency, mice were administered serial dilutions of TIS (1/2, 1/5, 1/10, 1/50, 1/500, and 1/1000) i.p. on days 0, 7, and 12. Tumor burden was evaluated on day 23 by measuring the total weight of excised peritoneal tumors. To evaluate the long-term protective effects of TIS, C57BL/6 mice were injected i.p. with TIS on days 0 and 7, followed by delayed tumor challenge with MC38 cells. Three experimental groups were established:

Analysis I: tumor challenge on day 37, followed by a third TIS injection on day 40; Analysis II: third TIS injection on day 94, followed by tumor challenge on day 97; Analysis III: tumor challenge on day 127 without additional TIS injection.

Tumor burden was assessed 14 days after tumor challenge by excision and weighing of peritoneal tumors.

### Survival, tumor rechallenge, and cross-species TIS transfer assays

For long-term survival analysis, C57BL/6 mice were treated with TIS and challenged i.p. with MC38 colon carcinoma cells (3 × 10⁵ cells/mouse) according to the experimental schedules described above. Mice were monitored for overall survival for up to 200 days (∼6 months). For tumor rechallenge, mice that survived the initial tumor challenge were i.p. re-injected with MC38 cells (3 × 10⁵ cells/mouse), together with age-matched naïve controls. Fourteen days after rechallenge, mice were euthanized, and tumor burden was evaluated by excising and weighing visible peritoneal tumor nodules. Peritoneal cells were harvested by lavage at the study endpoint, stained with fluorochrome-conjugated antibodies, and analyzed by flow cytometry. Gating was performed on live, singlet CD45⁺ cells to quantify major immune populations, including CD4⁺ and CD8⁺ T cells, CD19⁺ B cells, Siglec-F⁺ eosinophils, F4/80⁺ macrophages, and CD11c⁺ DCs. In the xenogeneic group (human-derived TIS into C57BL/6 mice), a subset of mice was i.p. challenged with MC38 cells (3 × 10⁵ cells/mouse) on day 60 (Figure 6B, Analysis IV). 14 days later, mice were euthanized, and tumor burden was assessed by excising and weighing visible peritoneal nodules.

### Allogeneic and xenogeneic TIS transfer for safety assessment

To evaluate the safety of TIS administration, C57BL/6 mice were injected i.p. with PBS, LPS, or TIS on days 0, 1, and 2, followed by repeat injections on days 7, 8, and 9. To test cross-species compatibility, TIS were prepared from C57BL/6 (C-TIS), Balb/c (B-TIS), or human CD3⁺ T cells (H-TIS) and transferred i.p. into recipient mice (C57BL/6 or Balb/c). Mice were monitored daily for survival and body weight changes for up to 60 days after the first injection. Body weight was recorded at each indicated time point, and percent change relative to baseline was calculated.

### In vivo cancer therapeutic model

To assess the therapeutic efficacy of TIS, C57BL/6 mice were inoculated i.p. with MC38 colon carcinoma cells (3 × 10⁵ cells/mouse; day 0). Beginning on day 3 after tumor inoculation, mice received intraperitoneal injections of PBS, LPS, or TIS on days 3, 5, 7, 9, and 11, corresponding to five every-other-day doses. On day 14, mice were euthanized, and tumor burden was evaluated by excising and weighing visible peritoneal tumor nodules. Peritoneal cells were harvested by lavage at the study endpoint, stained with fluorochrome-conjugated antibodies, and analyzed by flow cytometry. Gating was performed on live, singlet CD45⁺ cells to quantify major immune populations, including CD4⁺ and CD8⁺ T cells, CD19⁺ B cells, Siglec-F⁺ eosinophils, F4/80⁺ macrophages, and CD11c⁺ DCs.

### Macrophage and eosionophil depletion studies

To assess the role of macrophages in TIS-mediated anti-tumor effects, C57BL/6 mice were treated with clodronate liposomes (Liposoma BV, Amsterdam, Netherlands) according to the schedule shown in Figure 7A. TIS were administered i.p. as indicated, followed by MC38 tumor challenge (3 × 10⁵ cells/mouse). Fourteen days after tumor inoculation, mice were euthanized, and peritoneal immune cells and tumor nodules were collected. Tumor burden was quantified by weighing excised tumors, and immune profiling was performed by flow cytometry. For eosinophil depletion, mice were injected with anti–Siglec-F monoclonal antibody or rat anti-IgG2A Isotype Control antibody according to the schedule shown in Figure 7F. TIS treatment and MC38 tumor challenge were performed as above. On day 14 after tumor inoculation, peritoneal tumors were excised and weighed, and peritoneal immune composition was analyzed by flow cytometry.

### Immunophenotyping by flow cytometry

Isolated cells were first stained with a LIVE/DEAD viability dye (Thermo Fisher Scientific) to exclude non-viable cells. For surface marker analysis, cells were incubated with fluorochrome-conjugated antibodies for 1 h at 4 °C in FACS buffer (PBS containing 2% FBS and 2 mM EDTA). Cells were fixed using IC Fixation Buffer (BD Biosciences) according to the manufacturer’s protocols. Flow cytometry data were acquired on a BD LSR Fortessa™ (BD Biosciences) or CytoFLEX (Beckman Coulter) flow cytometer and analyzed using FlowJo software (Tree Star, Ashland, OR, USA).

### Scanning electron microscopy

For scanning electron microscopy (SEM), cells were fixed with 2.5% glutaraldehyde in PBS for 2 h, rinsed with PBS for 5 min, and post-fixed with 1% osmium tetroxide for 2 h. Samples were dehydrated through a graded ethanol series (30%, 50%, 70%, 90%, 100%) over 30 min, followed by drying in a critical point dryer (Leica). Specimens were sputter-coated with a 1–2 nm layer of gold–palladium and examined using a field-emission SEM (Hitachi, Tokyo, Japan).

### Statistical analysis

All data were analyzed using GraphPad Prism 8.0 (GraphPad Software, San Diego, CA, USA). Statistical differences between two groups were assessed using a two-tailed unpaired Student’s t-test. For comparisons involving three or more groups, one-way or two-way ANOVA with appropriate post hoc tests (Tukey’s or Bonferroni) was applied as indicated. Survival data were analyzed using the Kaplan–Meier method, and group comparisons were performed using the log-rank (Mantel–Cox) test. P values less than 0.05 were considered statistically significant. Exact n values and statistical test results are provided in the figure panels and corresponding legends. Statistical significance was defined as follows: P < 0.05 (*), P < 0.01 (**), P < 0.001 (***), P < 0.0001 (****), and n.s. (not significant). Data are presented as mean ± standard error of the mean (SEM) or standard deviation (SD), as indicated.

## Acknowledgements

This work was supported by the National Science Challenge Initiatives (RS-2024-00419699), the Mid-career Research Grant (RS2024-00341869), the Creative Research Initiative Program (2015R1A3A2066253), Sejong Science Fellowship Grant program (RS-2024-00340032), the Basic Science Program (2022R1A2C4002627) through National Research Foundation grants funded by the Ministry of Science and Information and Communication Technology and a grant of the Korea Health Technology R&D Project (RS-2024-00512909) through the Korea Health Industry Development Institute, funded by the Ministry of Health & Welfare, Republic of Korea. This work was supported in part by the National Cancer Center Grant (NCC2510661-1).

## Author contributions Statement

S.-K.K. and N.-Y.K. conceived the study. S.L., H.L., W.-C.S., J.-S.P., and H.-T.K. designed and performed the experiments. J.P., S.-J.L., Y.-J.S., and J.-H.K. performed microarray, bulk RNA-seq, and single-cell RNA-seq data analyses. H.-R.K. and C.-D.J. wrote and finalized the manuscript. All authors discussed the results and approved the final version of the manuscript.

## Supplemental Figure Legends

**Figure S1. Generation and purification of 4-TIS from activated CD4+ T cells**

(A) Confocal imaging of OT-II CD4⁺ T blasts stimulated on anti–CD3/CD28–coated surfaces. Cell body (CTV, blue), F-actin (TRITC-phalloidin, red), and TCRβ (Alexa Fluor 488, green). Internalized TCR signals (white arrows) and the release of TCR⁺ T cell immunosynaptosomes (TIS, boxed area) are shown. Middle: phase-contrast images of T cells incubated for 3 h with IgG- or anti-CD3/CD28–conjugated beads, showing selective adhesion to the latter. Right: confocal image of TCR⁺ TIS bound to beads. Schematic: bead-based activation and vortex-mediated release of TIS, followed by sucrose-gradient ultracentrifugation. Representative SEM images of purified TIS (18,000×). Western blot analysis of sucrose fractions probed with antibodies against TCRζ and ARRDC1.

(B) Quantification of TIS production from PMS-green–labeled OT-II CD4⁺ T cells. Left: representative flow cytometry plots (V-SSC vs PMS-green) of NT (no treatment) and iAb (anti-CD3/CD28–stimulated) conditions. Right: enumeration of TCRβ⁺ TIS particles over time under different stimulation conditions (NT, soluble Ab, peptide–MHC/ICAM-1).

(C) Functional stability of TIS. Peritoneal macrophages were stimulated with fresh (F-TIS), lyophilized (L-TIS), or frozen TIS (4-TIS). Nitrite concentrations (left) and TNF-α production (right) were quantified by the Griess assay and ELISA, respectively.

Bars represent mean ± SD of biological replicates. ns, not significant; *, p < 0.05; **, p < 0.01; ***, p < 0.001; ****, p < 0.0001.

**Figure S2. 4-TIS priming of GM-CSF–differentiated BMCs elicits adaptive-like cytokines, distinct from the inflammatory responses induced by LPS**

(A) Heatmap of cytokines measured by LEGENDplex cytometric bead array (CBA) from lysed CD4⁺ and CD8⁺ T cells under unstimulated (Uns-T) or stimulated (Stim-T) conditions, as well as from 4-TIS and 8-TIS. Samples were lysed, protein concentrations were normalized, and cytokine levels were quantified.

(B) Cytokine secretion by BMDCs after treatment with PBS, LPS, or TIS. Bar graphs show representative cytokines grouped by those predominantly induced by 4-TIS (top), shared between 4-TIS and LPS (middle), or preferentially induced by LPS (bottom).

(C) Cytokine secretion by BMDCs stimulated with 4-TIS generated from wild-type (WT) or IFN-γ–deficient (Ifng⁻/⁻) CD4⁺ T cells, showing dependence on IFN-γ.

(D) Cytokine secretion by BMDCs stimulated with 4-TIS in the presence or absence of blocking antibodies against TNF receptor (α-TNFR) or GM-CSF receptor (α-GM-CSFR). Bars represent mean ± SD. ns, not significant; *, p < 0.05; **, p < 0.01; ***, p < 0.001; ****, p < 0.0001.

**Figure S3. 4-TIS priming of peritoneal macrophages elicits adaptive-like cytokines, distinct from the inflammatory responses induced by LPS**

Bar graphs of cytokines secreted by peritoneal macrophages after treatment with PBS, LPS, or 4-TIS at the indicated concentrations. Cytokines are grouped into those preferentially induced by 4-TIS (top), comparably induced by 4-TIS and LPS (middle), or predominantly induced by LPS (bottom).

Bars represent mean ± SD. ns, not significant; *, p < 0.05; **, p < 0.01; ***, p < 0.001; ****, p < 0.0001.

**Figure S4. Systemic immune cell profiling after intraperitoneal injection of 4-TIS compared to conventional adjuvants.**

C57BL/6 mice were i.p. injected with PBS, LPS (25 μg), Poly(I:C) (50 μg), or 4-TIS on days 0, 3, and 6. On day 11, peritoneal cells were harvested and analyzed by flow cytometry.

Top row: Representative FACS plots and quantification of CD4⁺ and CD8⁺ T cells gated on CD3⁺ cells. CD4/CD8 ratio, absolute numbers of CD4⁺ and CD8⁺ T cells are shown. Middle row: Dendritic cells (DCs) were gated on CD45⁺MHCII⁺CD11c⁺. Frequencies and total numbers of DCs are presented.

Bottom row: B cells were identified as CD45⁺CD19⁺CD3⁻. CD19/CD3 ratio and total numbers of B cells are shown.

Data represent mean ± SEM (n = 5 per group). Statistical significance was determined by one-way ANOVA with Tukey’s post hoc test. *p < 0.05.

Bar graphs summarize total counts across time points (4 h, 1 d, 2 d, 8 d, and 11 d).

Bars represent mean ± SD. ns, not significant; *, p < 0.05; **, p < 0.01; ***, p < 0.001; ****, p < 0.0001.

**Figure S5. 4-TIS is preferentially captured by professional antigen-presenting cells in vivo**

(A) Flow cytometry of peritoneal cells at indicated time points after i.p. injection of PMS-green–labeled 4-TIS. CD45⁺ cells were gated and analyzed for uptake within macrophages (F4/80⁺), DCs (CD11c⁺), B cells (CD19⁺), eosinophils (Siglec-F⁺), and T cells (CD3⁺). Representative plots show PMS-green⁺ events over 0.5–72 h. Blue arrows indicate PMS-green⁺ populations.

(B) Quantification of PMS-green⁺ populations over time, shown as percentages of each lineage. Bars represent mean ± SD. ns, not significant; *, p < 0.05; **, p < 0.01; ***, p < 0.001; ****, p < 0.0001.

**Figure S6. Unlike LPS, 4-TIS induces delayed but broad immune cell recruitment with adaptive immune traits in the skin**

(A) Experimental scheme for intradermal (i.d.) injection of 4-TIS. Skin was collected at the indicated time points for histology and immune profiling. Representative H&E-stained skin sections are shown at days 1, 2, 8, and 11 after i.d. injection of PBS, LPS (25 μg), Poly(I:C) (50 μg), or 4-TIS. Pie charts depict the relative proportions of macrophages, lymphocytes, neutrophils, and eosinophils. Insets highlight representative lesions: neutrophil-dominant infiltration after LPS (day 2, red box) and eosinophil-rich infiltration after 4-TIS (day 8, blue box).

(B) Quantification of skin thickness (left) and infiltrating immune cells per high-power field (HPF, 400×; right) over the indicated time course.

(C) Cytokine and chemokine measurements after i.d. PBS, LPS (25 μg), Poly(I:C) (50 μg), or 4-TIS. Top: levels in skin tissue lysates; bottom: corresponding serum levels at days 1, 2, 8, and 11.

Bars show mean ± SD. ns, not significant; *, p < 0.05; **, p < 0.01; ***, p < 0.001; ****, p < 0.0001.

**Figure S7. 4-TIS increases ECAR while maintaining OCR in peritoneal B cells.**

B cells were isolated from peritoneal lavage and subjected to Seahorse extracellular flux analysis. Shown are representative traces of OCR (left) and ECAR (right) over time following sequential injections of mitochondrial stress test compounds (oligomycin, FCCP, and rotenone/antimycin A). B cells from mice treated with PBS, LPS, or 4-TIS were compared. Data represent mean ± SEM. *, p < 0.05; **, p < 0.01; ***, p < 0.001 (vs. PBS).

**Figure S8. Canonical marker gene expression defines peritoneal immune cell clusters in scRNA-seq**

Dot plot showing the expression of canonical marker genes used for cell type annotation across clusters 0–19 from scRNA-seq of peritoneal cells. Marker sets were adapted from Zhao et al. (Commun Biol 5, 1225, 2022). Dot color represents mean expression level, and dot size reflects the fraction of cells expressing the gene within each cluster.

**Figure S9. TIS-specific gene expression and transcriptional programs in proliferating macrophages, early memory B cells and DCs**

(A) UMAP of macrophages colored by the TIS-signature module score.

(B) Bar plot of GO biological processes enriched among genes upregulated in 4-TIS–induced proliferating macrophages.

(C) UMAP of early memory B cells from PBS, LPS, and 4-TIS, colored by treatment.

(D) Heatmap of early memory B cells DEGs across PBS, LPS, and 4-TIS (row-wise z-scores).

(E) Bar plot of GO biological processes enriched among 4-TIS-upregulated genes in early memory B cells.

(F) UMAP of DCs from PBS, LPS, and 4-TIS, colored by treatment.

(G) Heatmap of macrophage proliferation related genes evaluated in DCs (row-wise z-scores).

(H) Bar plot of GO biological processes enriched among 4-TIS-or LPS-upregulated genes in DCs.

**Figure S10. Validation of eosinophil and T cell sorting by lineage-specific marker expression**

(A) Bar graph showing RT-qPCR mRNA expression of *Cd3, Gata2, Ccr3, Siglecf, Epx, and Prg3* in sorted T cells (Tc) and eosinophils (Eos), normalized to *Gapdh*.

**Figure S11. 4-TIS-induced alterations in peritoneal immune cell populations with or without MC38 cell inoculation**

(A) Experiment design as in Figure 5A. Top: treatment schemes without MC38 inoculation; representative flow cytometry plots (FSC/SSC) and pie charts showing peritoneal immune cell composition (CD45⁺ gate) at day 23. Bottom: corresponding FSC/SSC plots and immune-cell composition for the Figure 5A treatment groups with MC38 inoculation.

**Figure S12. 4-TIS exhibits both long-lasting preventive and therapeutic anti-tumor effects**

(A) Schematic of the in vivo treatment timeline for the staggered booster experiment.

(B–E) Immune profiling of peritoneal cells at analysis (I–IV). Representative flow cytometry plots (FSC/SSC, CD45⁺ gate), pie charts of immune-cell composition, and bar graphs of total peritoneal cells and immune subsets (CD8⁺ T cells, CD4⁺ T cells, CD19⁺ B cells, eosinophils, neutrophils, DCs, macrophages, SPMs, and LPMs).

Bars represent mean ± SEM. ns, not significant; *, p < 0.05; **, p < 0.01; ***, p < 0.001; ****, p < 0.0001.

**Figure S13. 4-TIS reduces B16F10 lung metastasis and modulates systemic immunity**

(A) Schematic of the in vivo treatment timeline for the B16F10 intravenous (i.v.) lung metastasis model.

(B) Representative photographs of local skin lesions at the injection site after intradermal (i.d.) administration of PBS, LPS, or 4-TIS.

(C) Representative images of lungs harvested 21 days after the first injection, with bar graph quantification of metastatic B16F10 foci.

(D) Immune cell counts in draining lymph nodes (dLNs), including CD8⁺ T cells, CD4⁺ T cells, CD19⁺ B cells, CD4⁺ CXCR5⁺ T cells, and CD44⁺ T cells.

(E) Serum cytokine levels (IL-2 and IFN-γ) measured at day 21.

Bars represent mean ± SD. ns, not significant; *, p < 0.05; **, p < 0.01; ***, p < 0.001; ****, p < 0.0001.

**Table S1. List of primers used for RT-qPCR analysis**

