## Supplementary figures and images for "T-Cell Synaptosomes Orchestrate Long-Term Anti-Tumor Immunity via Proliferative and Metabolic Reprogramming"

### Supplemental Figure 1

Figure S1. Generation and purification of 4-TIS from activated CD4⁺ T cells

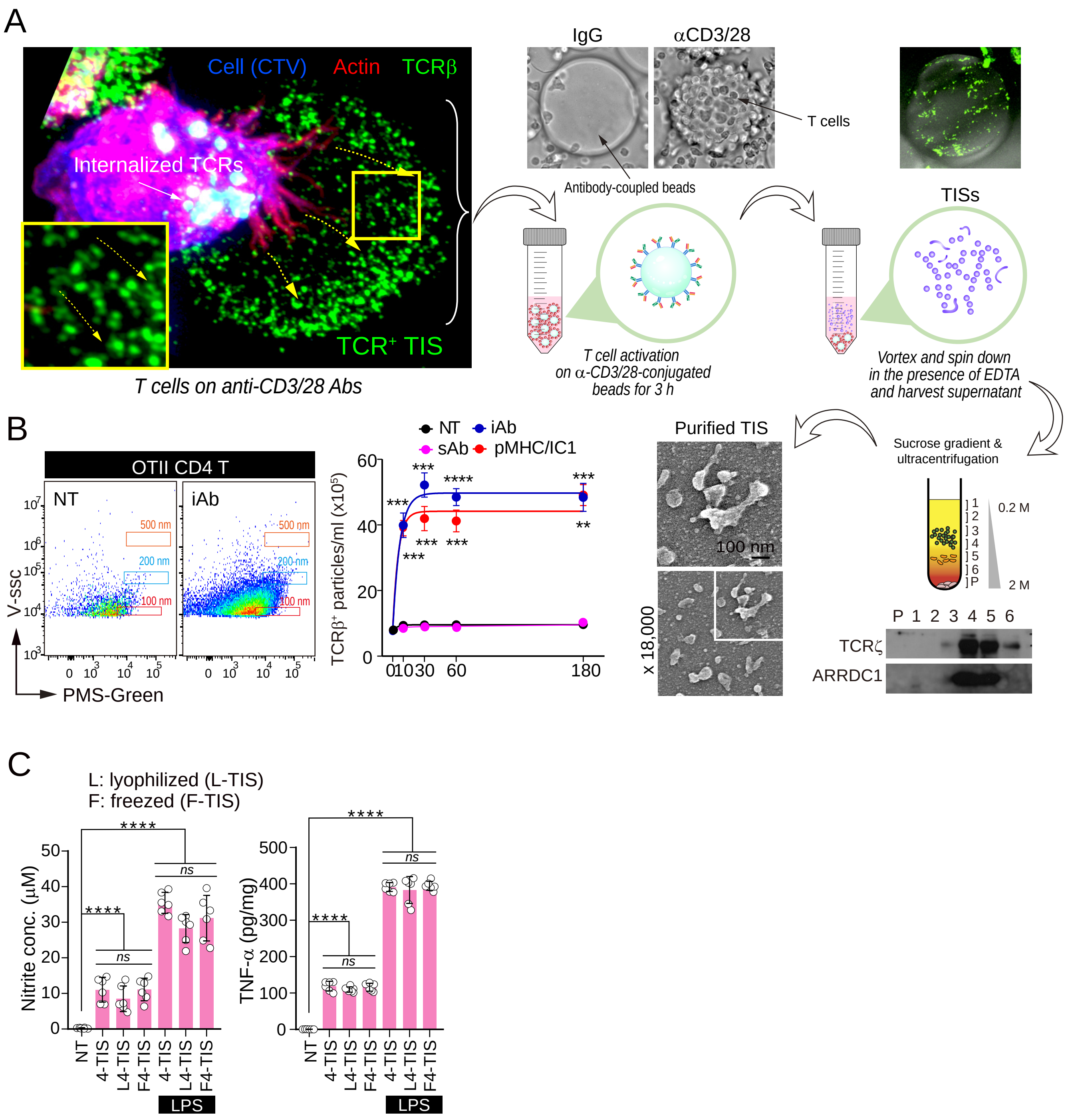

### Supplemental Figure 5

Figure S5. TIS is preferentially captured by the professional antigen-presenting cells *in vivo*

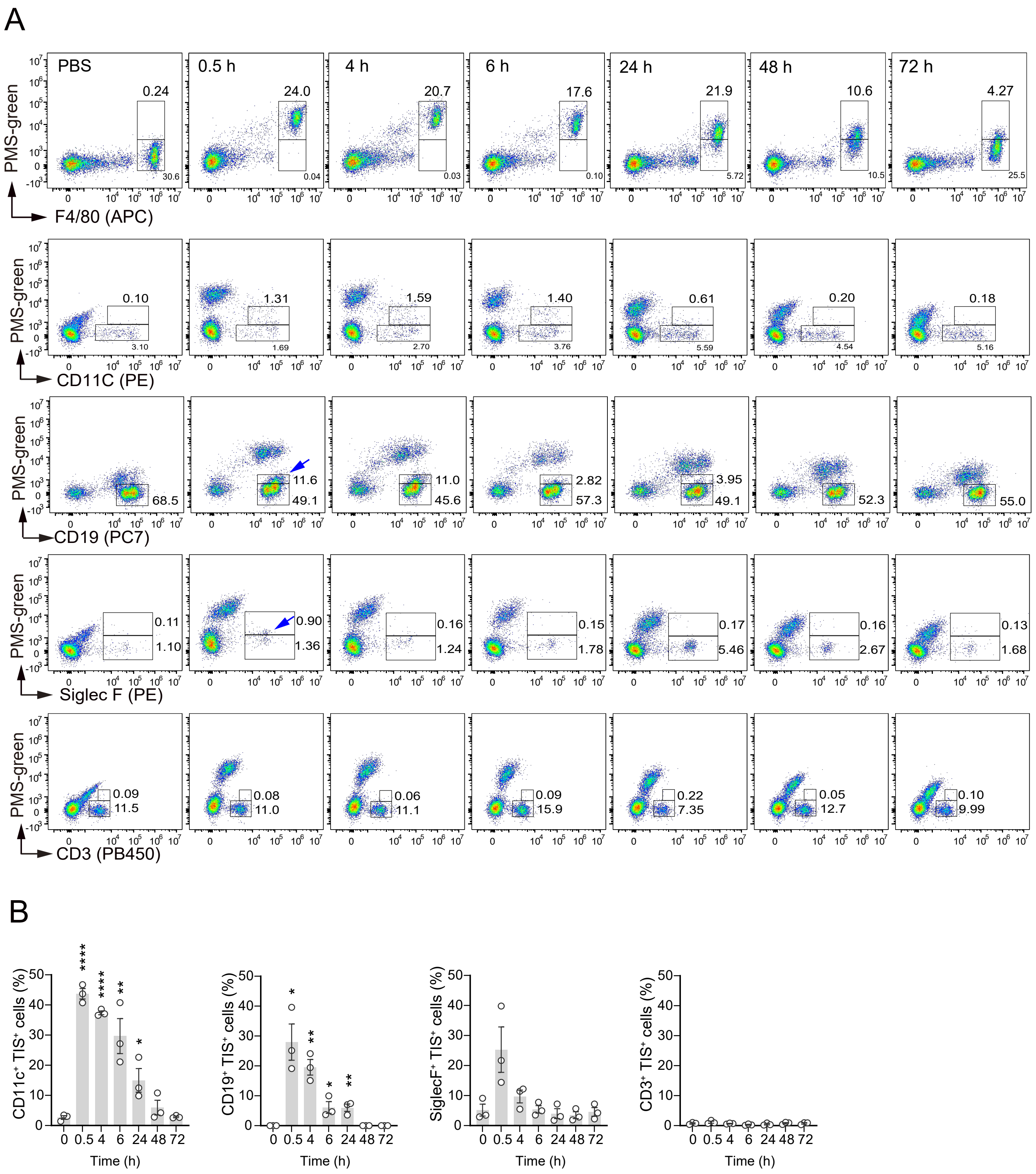

### Supplemental Figure 7

Figure S7. 4-TIS increases ECAR while maintaining OCR in peritoneal B cells

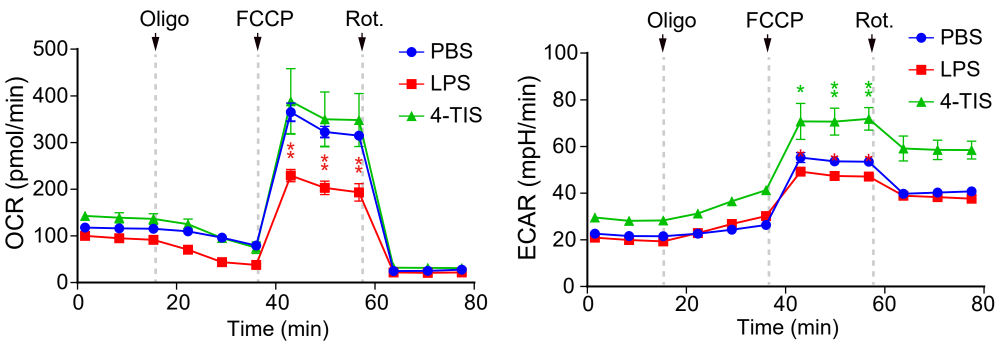

### Supplemental Figure 10

Figure S10. Validation of eosinophil and T cell sorting by lineage-specific marker expression

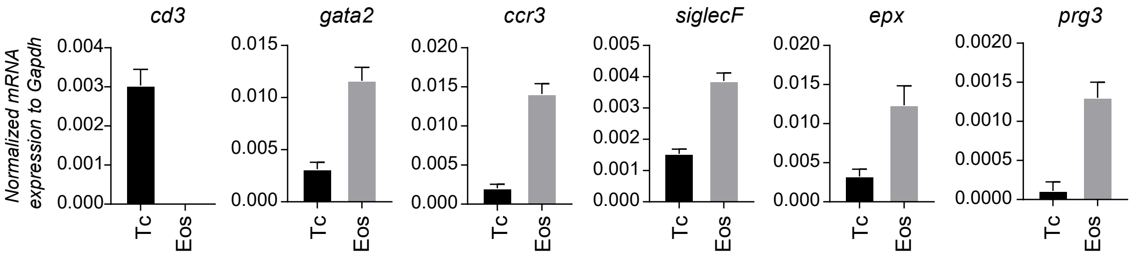

### Supplemental Figure 13

Figure S13. TIS reduces B16F10 lung metastasis and modulates systemic immunity

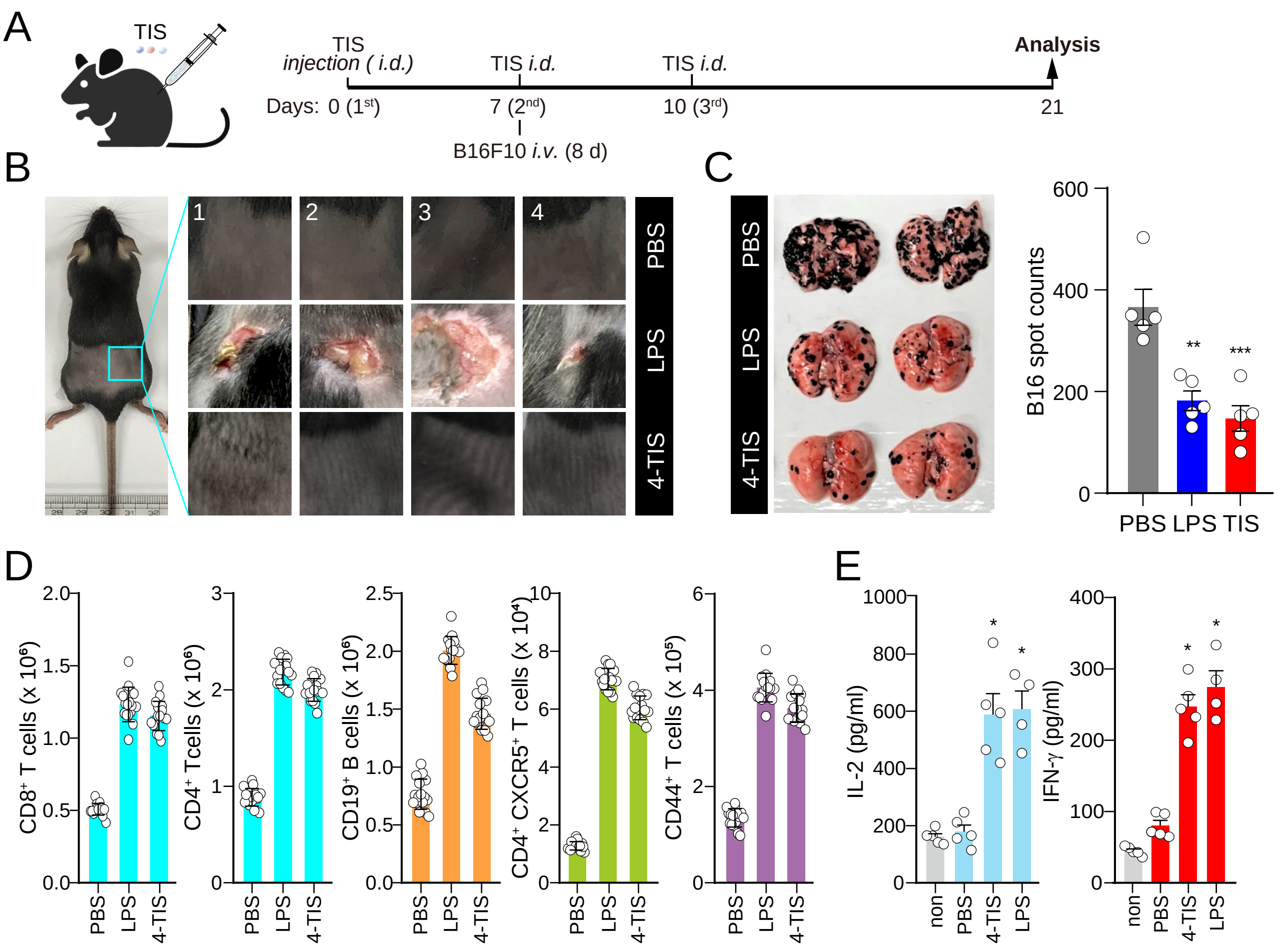
