## Supplemental Figure 2 for "T-Cell Synaptosomes Orchestrate Long-Term Anti-Tumor Immunity via Proliferative and Metabolic Reprogramming"

Figure S2. 4-TIS priming of GM-CSF-differentiated BMCs elicits adaptive-like cytokines, distinct from the inflammatory responses induced by LPS

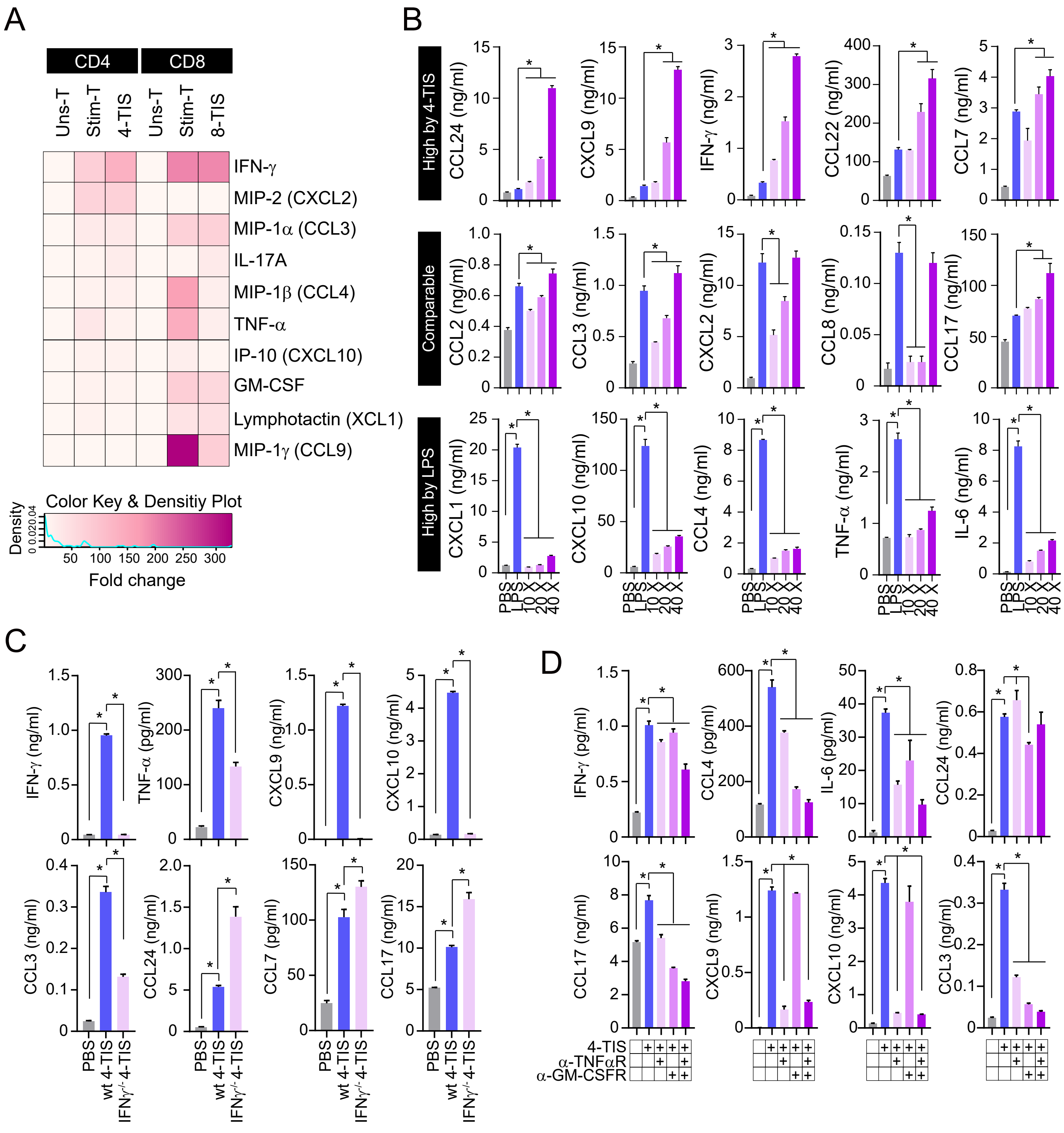
