## Supplemental Figure 3 for "T-Cell Synaptosomes Orchestrate Long-Term Anti-Tumor Immunity via Proliferative and Metabolic Reprogramming"

Figure S3. 4-TIS priming of peritoneal macrophages elicits adaptive-like cytokines, distinct from the inflammatory responses induced by LPS

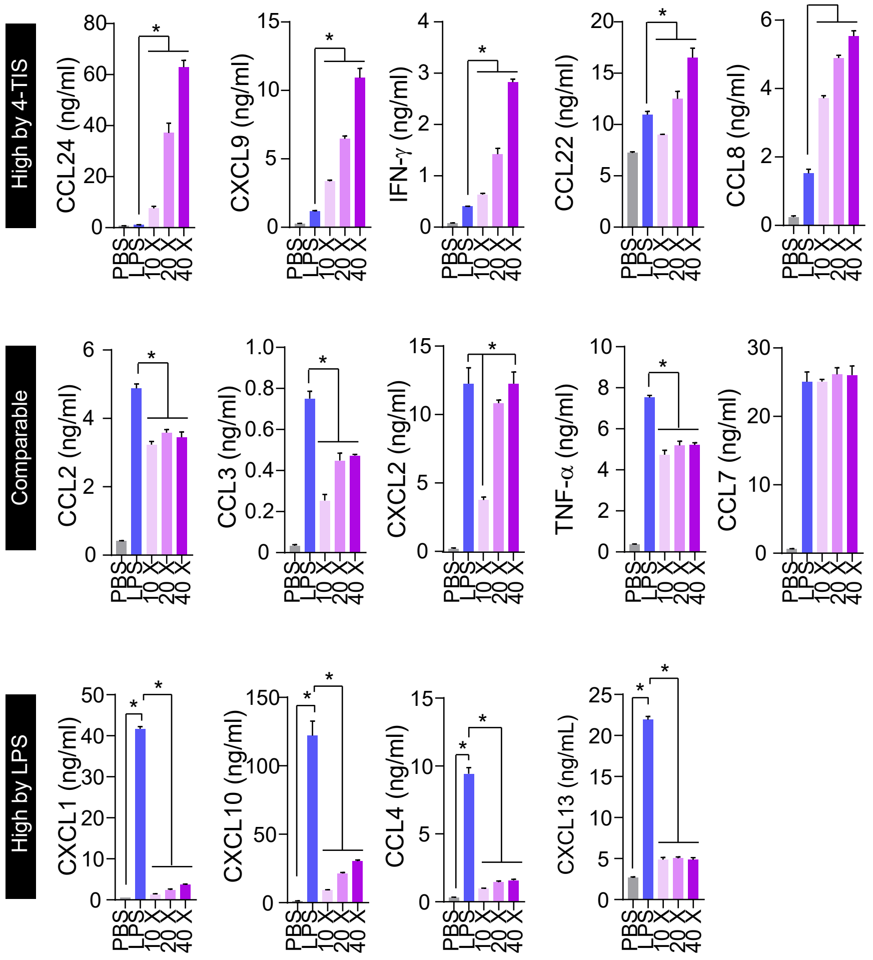
