## Supplemental Figure 4 for "T-Cell Synaptosomes Orchestrate Long-Term Anti-Tumor Immunity via Proliferative and Metabolic Reprogramming"

Figure S4. 4-TIS promotes delayed and broad immune cell recruitment with adaptive immune features unlike LPS

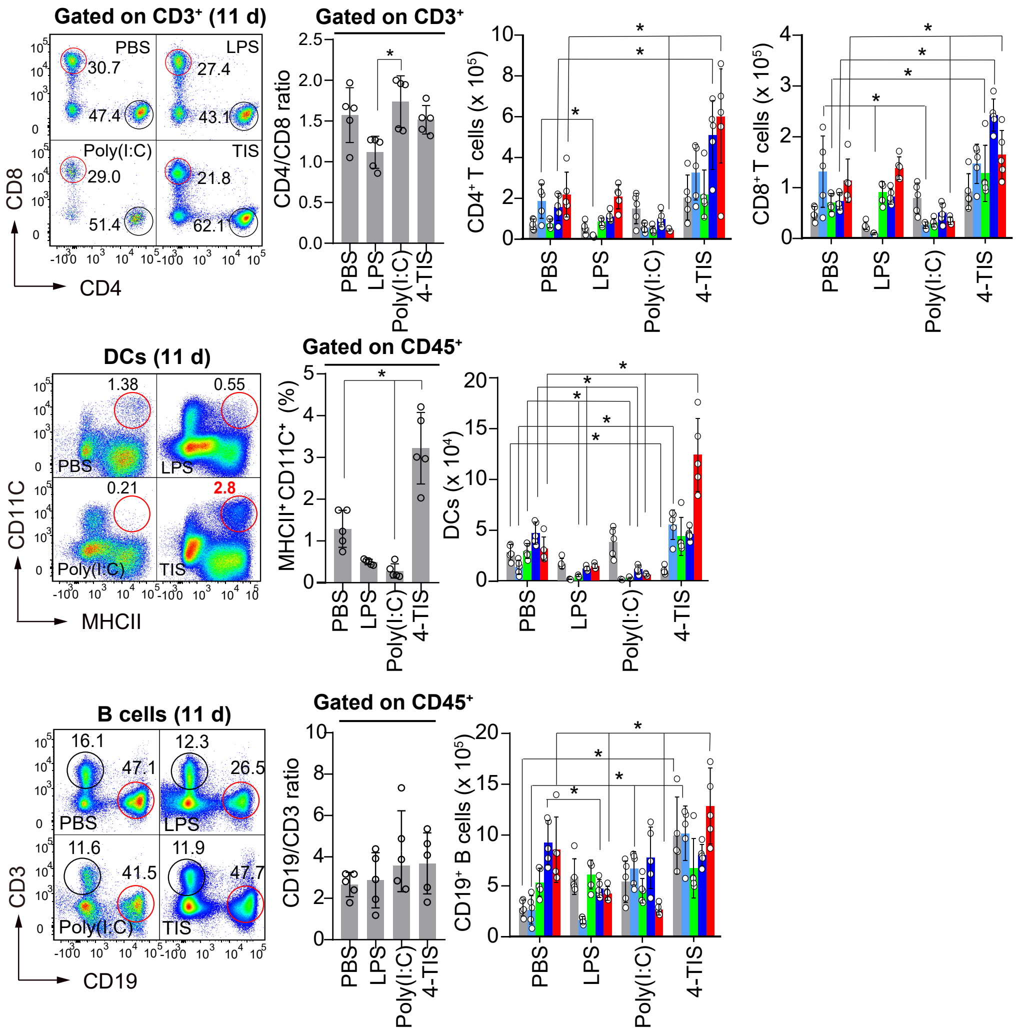
