## Supplemental Figure 6 for "T-Cell Synaptosomes Orchestrate Long-Term Anti-Tumor Immunity via Proliferative and Metabolic Reprogramming"

Figure S6. Unlike LPS, 4-TIS induces delayed but broad immune cell recruitment with adaptive immune traits in the skin

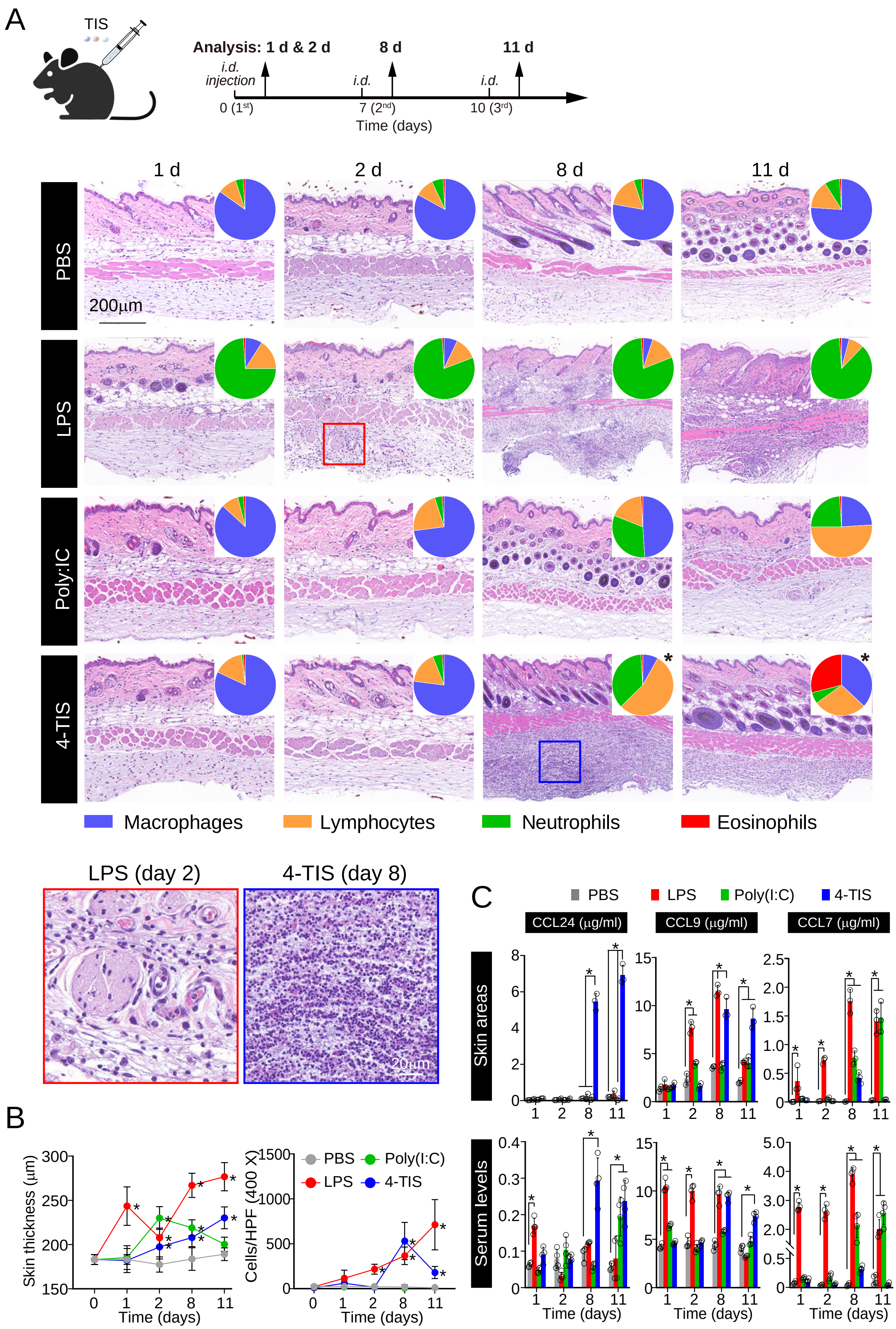
