## Supplemental Figure 8 for "T-Cell Synaptosomes Orchestrate Long-Term Anti-Tumor Immunity via Proliferative and Metabolic Reprogramming"

Figure S8. Canonical marker gene expression defines peritoneal immune cell clusters in scRNA-seq

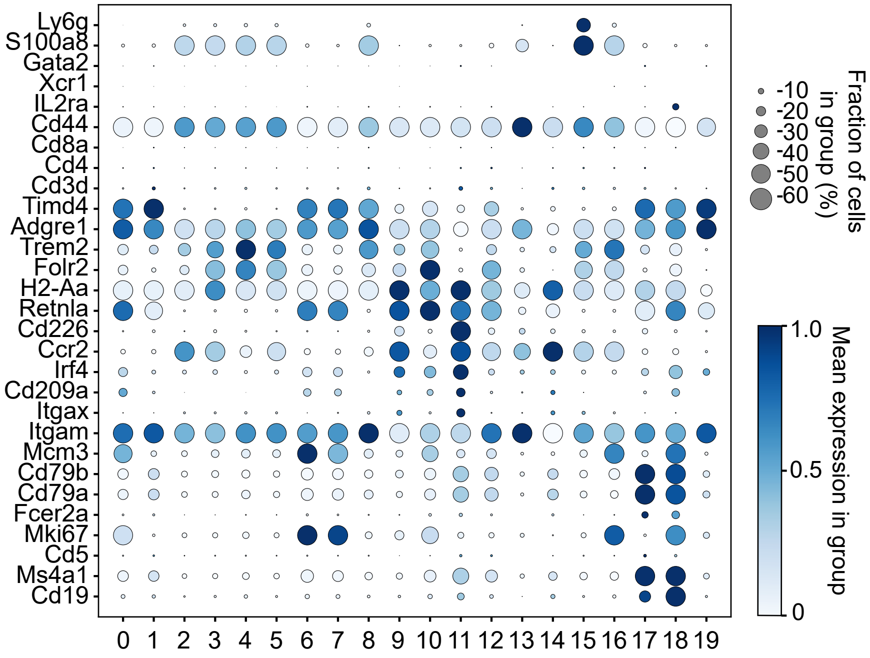
