## Supplemental Figure 9 for "T-Cell Synaptosomes Orchestrate Long-Term Anti-Tumor Immunity via Proliferative and Metabolic Reprogramming"

Figure S9. TIS-specific gene expression and transcriptional programs in macrophages, B cells and DCs

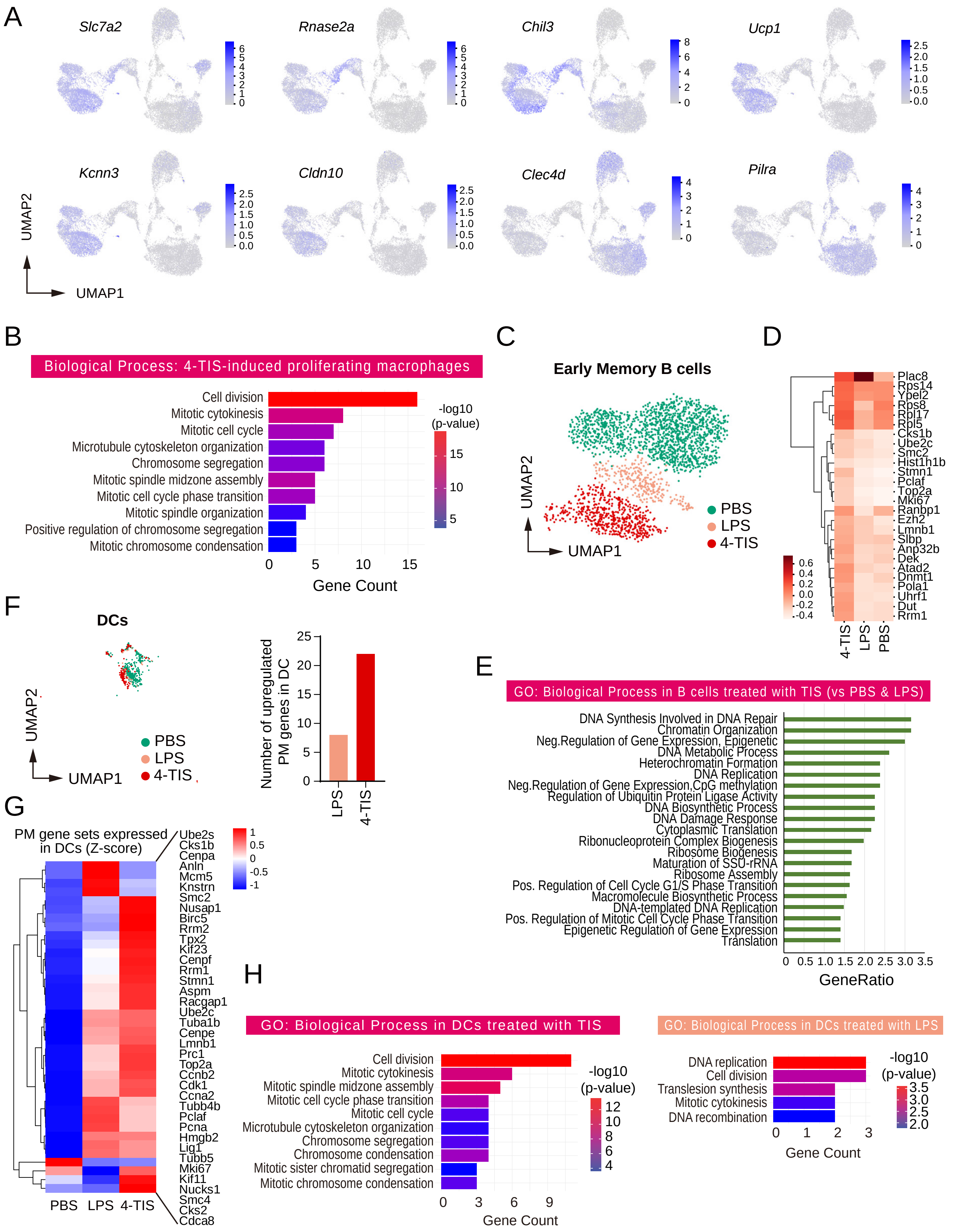
