## Supplemental Figure 11 for "T-Cell Synaptosomes Orchestrate Long-Term Anti-Tumor Immunity via Proliferative and Metabolic Reprogramming"

Figure S11. 4-TIS-induced alterations in peritoneal immune cell populations with or without MC38 cell inoculation

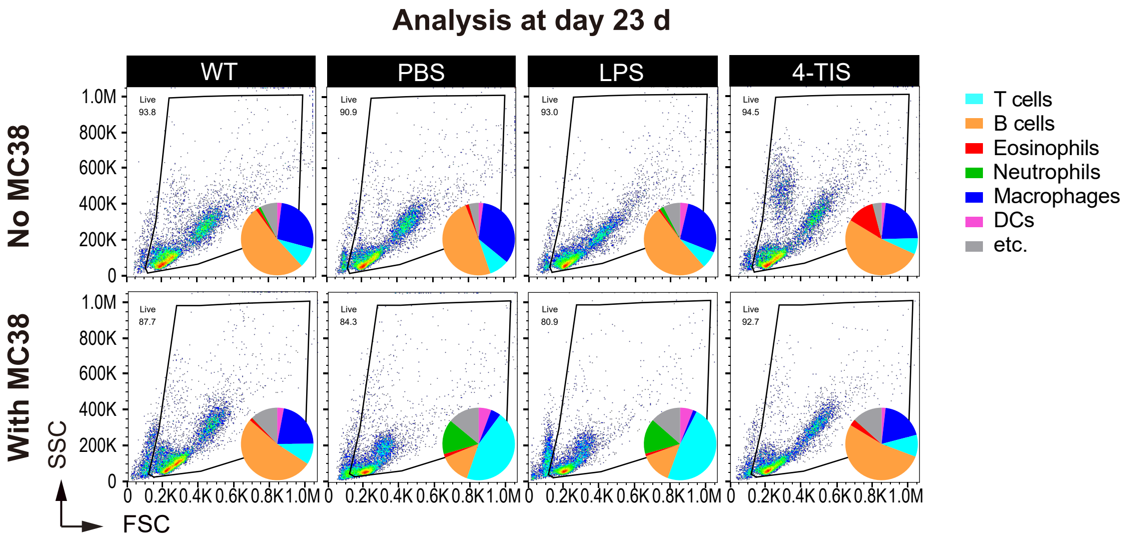
