## Supplemental Figure 12 for "T-Cell Synaptosomes Orchestrate Long-Term Anti-Tumor Immunity via Proliferative and Metabolic Reprogramming"

### Figure S12. TIS exhibits both long-lasting preventive and therapeutic anti-tumor effects

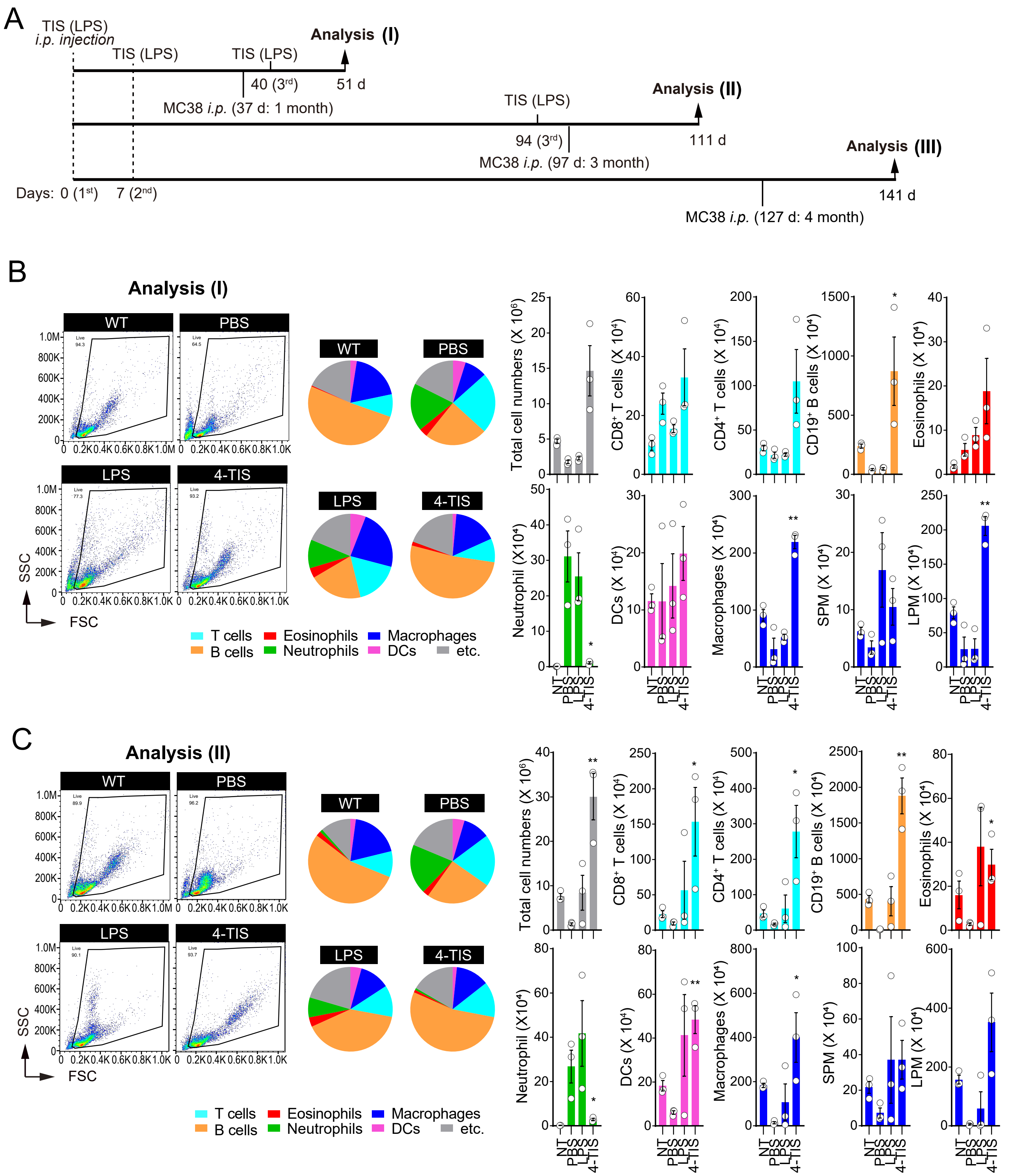

Figure S12. TIS exhibits both long-lasting preventive and therapeutic anti-tumor effects (continue)

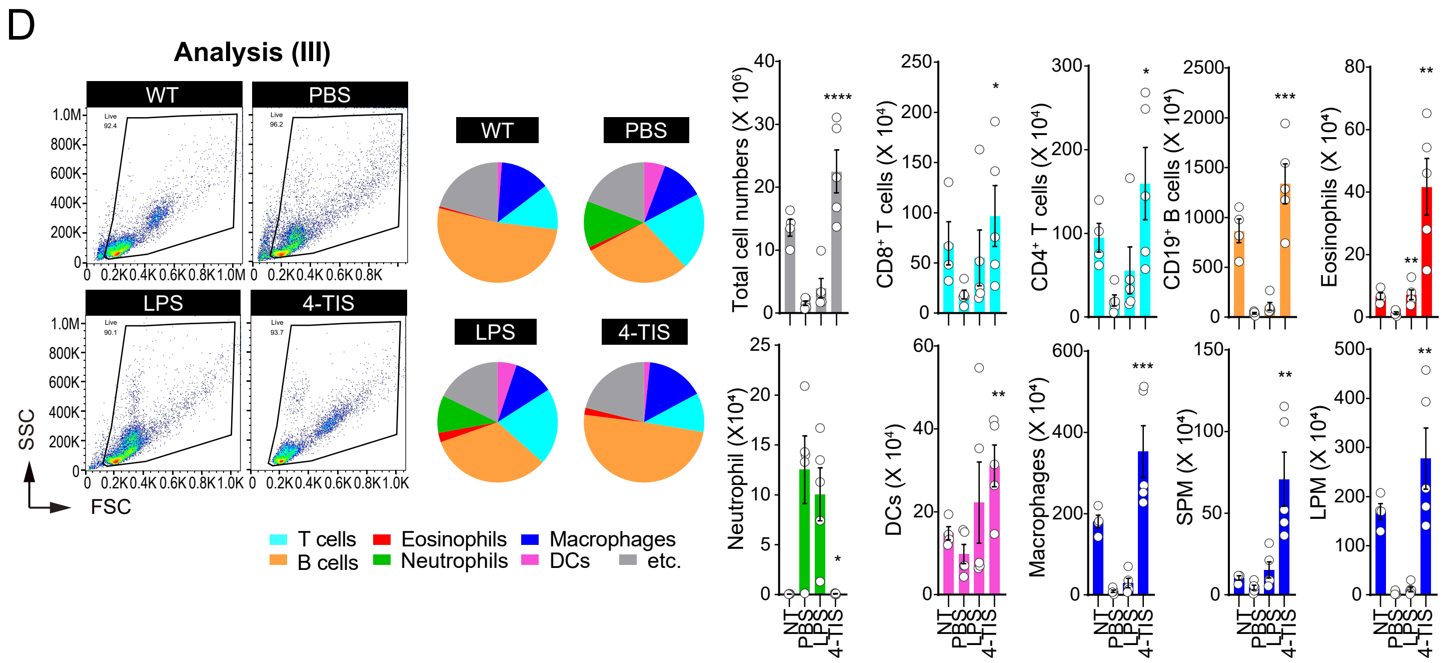


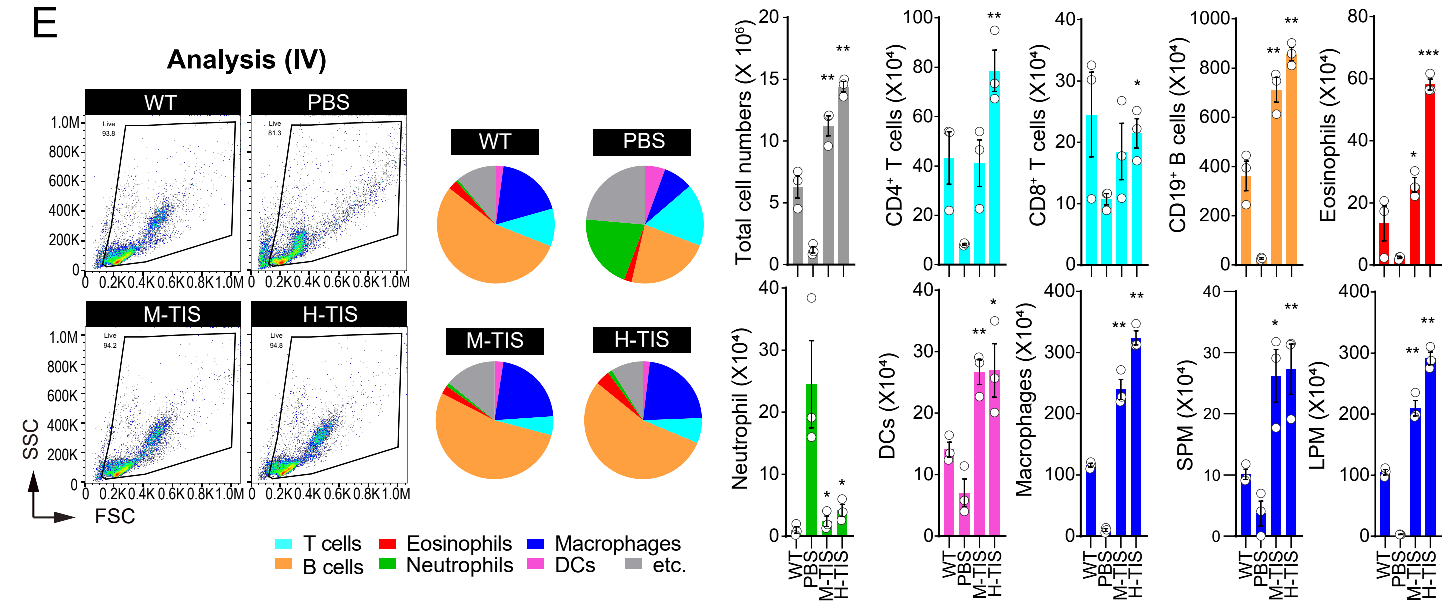
