## Supplemental Table 1 for "T-Cell Synaptosomes Orchestrate Long-Term Anti-Tumor Immunity via Proliferative and Metabolic Reprogramming"

Table S1. List of primers used for RT-qPCR analysis

| | | |
|---|---|---|
| *gata1* | Forward Primer | TGGGGACCTCAGAACCCTTG |
| | Reverse Primer | GGCTGCATTTGGGGAAGTG |
| *gata2* | Forward Primer | CACCCCGCCGTATTGAATG |
| | Reverse Primer | CCTGCGAGTCGAGATGGTTG |
| *Siglec-f* | Forward Primer | GCTGGTCTTATGGCCTTGCTT |
| | Reverse Primer | AGGCTCATCCTTTTGGGTGTC |
| *prg2* | Forward Primer | GTCTCAGGTCAGGATGTGACA |
| | Reverse Primer | GCGGACTGGATTCCGAAGTT |
| *prg3* | Forward Primer | ACAGCCCCTGATCCTGTCC |
| | Reverse Primer | CTTGCTCCCTCGAACCATCTG |
| *epx* | Forward Primer | CTCACCCAACACGCTGAAG |
| | Reverse Primer | TTTTCTGTGTGTGATTGTAGGCA |
| *ccr3* | Forward Primer | TCAACTTGGCAATTTCTGACCT |
| | Reverse Primer | CAGCATGGACGATAGCCAGG |
| *cd3e* | Forward Primer | ATGCGGTGGAACACTTTCTGG |
| | Reverse Primer | GCACGTCAACTCTACACTGGT |
| *msdha* | Forward Primer | TCGACAGGGGAATGGTTTGG |
| | Reverse Primer | TCATACTCATCGACCCGCAC |
| *ccna2* | Forward Primer | GAGGTCCTAACGCTCCCATC |
| | Reverse Primer | TCTGGCCTACATGTCCTCTG |
| *mki67* | Forward Primer | ACCATCATTGACCGCTCCTTT |
| | Reverse Primer | TTGACCTTCCCCATCAGGGT |
| *ccl24* | Forward Primer | CCTGAACTTGGACATAGGGG |
| | Reverse Primer | TGATGAAGCCTTTGAGCCACA |
| *ccl4* | Forward Primer | TGTGCAAACCTAACCCCGAG |
| | Reverse Primer | CCATTGGTGCTGAGAACCCT |
| *itgb7* | Forward Primer | TTGGATGATGGCTGGTGCAA |
| | Reverse Primer | CTAGTCCCACTGCCACGATG |
| *ifna1* | Forward Primer | AGGACTTTGGATTCCCGCAG |
| | Reverse Primer | TCATTGAGCTGCTGGTGGAG |
